# Emergent dynamics of cellular decision making in multi-node mutually repressive regulatory networks

**DOI:** 10.1101/2025.03.06.641782

**Authors:** BV Harshavardhan, Hanuma Sai Billakurthi, Sarah Adigwe, Kishore Hari, Herbert Levine, Tomas Gedeon, Mohit Kumar Jolly

## Abstract

Stem cell differentiation during development is governed by the dynamics of the underlying gene regulatory networks (GRNs). Mutually inhibiting nodes/collection of nodes encompass the GRNs that govern differentiation to two distinct fates. But the properties of GRNs that can allow differentiation into n-terminal phenotypes are not understood. In this study, we examine toggle-n networks, encompassing mutual inhibitions among multiple transcription factors, to derive generalized conclusions regarding the dynamics underlying differentiation into n-terminal phenotypes. We show that in steady-state distributions of gene expression, multiple cell state-specific transcription factors are co-expressed, indicating an obligatory multi-step process of multi-lineage differentiation. Furthermore, we show that cytokine signaling and specific asymmetry of regulatory links can lead to directed differentiation towards a particular cell state. Our findings provide valuable insights into the mechanistic aspects of directed differentiation of stem cells.

## 1 Introduction

Elucidating the design principles of cell fate decisions has profound implications for understanding developmental biology, regenerative medicine, and synthetic biology (Qian et al., 2018; Sáez et al., 2022a; Zhou & Huang, 2011). The process of cell fate determination can be conceptualized through the metaphor of Waddington’s landscape, illustrating the myriad trajectories a cell can undertake during differentiation. Each valley in this landscape represents a terminally differentiated phenotype defined by specific gene expression patterns and underlying epigenetic structures. Each branch represents the event of differentiation of a precursor or progenitor cell to the sister lineages to which it could give rise (usually two or three). These trajectories are finely regulated by the dynamics of gene regulatory networks (GRNs), which orchestrate gene expression patterns and shape the landscape, thus guiding cells toward specific fates (Furusawa & Kaneko, 2012; Sáez et al., 2022b). These GRNs often exhibit multistability, which allows them to adopt any of these distinct stable states, where each corresponds to different cellular phenotypes (Ghaffarizadeh et al., 2014; Guantes & Poyatos, 2008). Central to this concept is the notion of a (terminally) differentiated cell expressing cell state-specific transcription factors (TFs) at high levels. This expression pattern highlights the precision in the commitment of a cell to a particular lineage, marked predominantly by expressing the specific TFs. It should be noted that (terminally) differentiated cells could show plasticity and switch to another cell state, which may not be apparent from the term “terminal” (Alvarado & Yamanaka, 2014). Cell state-specific TFs for a particular cell type tend to be co-expressed for a particular cell state, and the TFs specific to other cell states are excluded. These co-expression patterns could be a possible outcome of the existence of teams of nodes such that TFs within the same team activate each other while inhibiting TFs that belong to other teams. This mutual activation and inhibition ensure the stability of the cell state by reinforcing the expression of the TFs within the same state while suppressing the expression of those associated with alternate states (Hari et al., 2022b).

Cell-fate choices between two and three possible states are well studied using a toggle switch (GRN featuring mutual inhibition between two transcription factors (TFs)) and a toggle triad (GRN featuring mutual inhibition between three TFs) through ordinary differential equation (ODE)-based formalism (Duddu et al., 2020; Gardner et al., 2000; Ma et al., 2006; Tian & Burrage, 2006; Yang et al., 2021) and boolean approaches (Hari et al., 2022a; Masashi et al., 2017; Zhou et al., 2016). Also, the role of extracellular signaling in directing differentiation trajectories has been elucidated using dynamical systems approaches, particularly for a toggle switch (Pezzotta & Briscoe, 2023; Wang et al., 2022). However, a stem cell can differentiate into more than three cell fates; for example, a hematopoietic stem cell can differentiate into various terminal cell types, including erythrocytes, basophils, eosinophils, neutrophils, megakaryocytes, monocytes and lymphocytes (Laurenti & Göttgens, 2018; Zhou & Huang, 2011). The dynamics of networks that may enable more than two or three possible cell states need to be analyzed further.

Recent studies have ventured into larger GRNs with mutual inhibition, the toggle tetrahedron (featuring mutual inhibitions between four TFs), where single positive states corresponding to the terminally differentiated phenotypes were not the predominant ones, necessitating additional mechanisms for complete differentiation (Duddu et al., 2024; Hong et al., 2015). Whether this pattern is an anomaly or a consistent trend with larger networks requires further investigation. In this study, we extend these findings by examining networks featuring mutual inhibitions, specifically toggle-n (Tn) networks, where n denotes the number of nodes, to derive generalized conclusions regarding the dynamics underlying differentiation into n terminal phenotypes. We analyze such networks with a varying number of nodes, ranging from 2 to 8 (Figures 1A and 1B), utilizing Boolean simulations employing an Ising formalism (Font-Clos et al., 2018). To complement the choice of a particular update formalism, we consider a collection of all monotone Boolean functions that are compatible with the given network, and describe the collection of steady states in this ensemble of functions.

**Figure 1:**
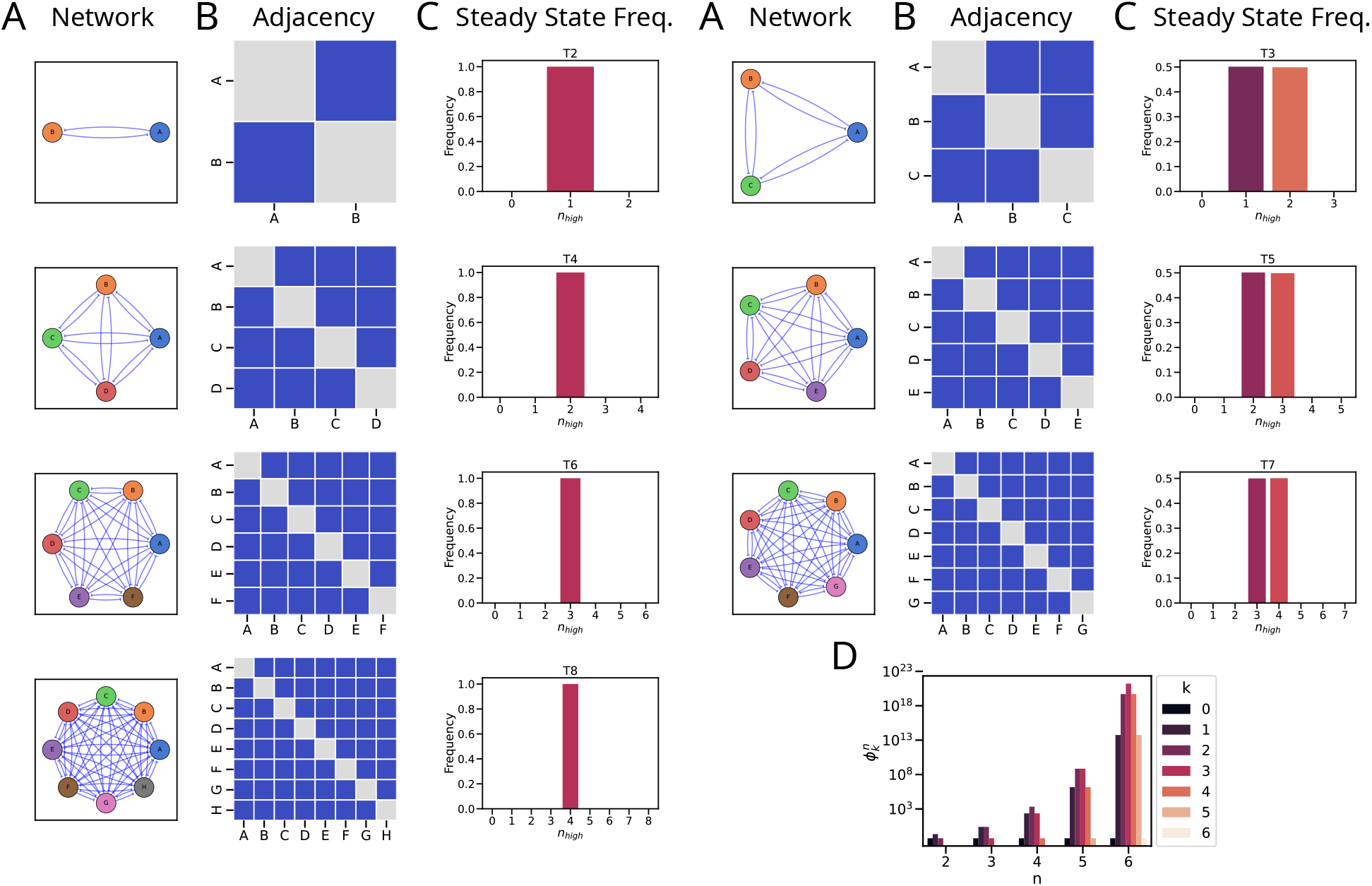
Schematic representation of the toggle-n networks and their steady states: A: Graphical network representation illustrating the nodes and edges representing TFs and interactions respectively. B: Adjacency matrix representation with the rows/columns and elements representing the interaction, respectively. Red/+1, Blue/-1, and Grey/0 represent activatory, inhibitory, and no interactions. C: Steady state frequency distribution of _*nhigh*_ states. Left: Even networks, Right: Odd networks. D: Number of monotone Boolean functions supporting steady states of *k*-high states for networks with *n* nodes.

We find that in steady-state distributions of gene expression, TFs are co-expressed from state-specific TFs from multiple states, indicating the need for a multi-step process of multi-lineage differentiation. These traits hold true even under random variations in regulatory link strengths and influence of other genes. Additionally, we show that cytokine signaling and specific asymmetry of regulatory links can lead to directed differentiation towards a particular cell state. Overall, our results suggest that while toggle-n networks play a pivotal role, they necessitate additional mechanisms to effectively orchestrate the trajectories of (terminal) cell differentiation.

## 2 Results

### 2.1 Mutually repressive regulatory networks allow the cell to bifurcate into pre-cursor lineages

In our investigation, we analyzed the steady-state distributions arising from Toggle-n networks using asynchronous Boolean simulations with the Ising formalism; details are presented in the Methods section. We observed distinct frequency patterns of *n*_*high*_, i.e., the number of nodes/TFs that are ON (high) in individual steady states for these different-sized networks. For the networks with two (T2), four (T4), six (T6), and eight (T8) nodes, we obtained 1 high, 2 high, 3 high, and 4 high states, respectively. In other words, for a T6 network (i.e., six nodes mutually inhibiting one another), the steady state(s) show any three out of six nodes being ON (high) and the remaining three being OFF (low). The pattern for networks with three (T3), five (T5), and seven (T7) nodes have the frequencies shared between 1 high - 2 high, 2 high - 3 high, and 3 high - 4 high, respectively (Figure 1C). In other words, for a T5 network (i.e., five nodes mutually inhibiting one another), the steady state(s) show any two or three out of five nodes being ON (high) and the remaining three or two being OFF (low). Since these networks are symmetric, the frequencies of the individual states with the same *n*_*high*_ are equal. Furthermore, the frequency of states with *n*_*high*_ equals the frequency of states with 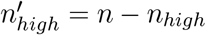.

We confirm these numerical results by analyzing the convergence properties of Ising update function dynamics. Based on ideas from Font-Clos et al., 2018, we show that the states with configuration *k* ≈ *n/*2 (where *k* represents the number of high nodes) exhibit the lowest energy within the system. Furthermore, the Ising update ensures a strict decrease in the energy function for any state distinct from the states with the lowest energy. As a consequence, these states with *k* ≈ *n/*2 act as attractors in the Ising update dynamics. These findings are summarized in the following Theorem, whose detailed proof can be found in Section S.1

**Theorem 1**. *Ising model update for network Tn for any n* ≥ 2, *applied to any initial condition will converge to a steady state with k high states, where*

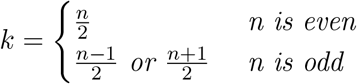

To investigate whether the choice of the Ising update function or the repressive structure of the networks is responsible for the distribution of the steady states, we explicitly constructed the set of all monotone Boolean functions that are compatible with each of the networks Tn for *n* = 2, 3, 4, 5, 6. We did not compute for *n* = 7, 8 as the number of monotone Boolean functions of *s* variables is the Dedekind number *D*(*s*) that grows extremely quickly and is only known for *s* ≤ 9 (Fidytek et al., 2001; Jäkel, 2023). We then computed the number of monotone Boolean functions 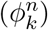 that support steady states with *k* = 0, 1, …, *n* high states in network *Tn* (Figure 1D). These numbers are listed in the ascending order below

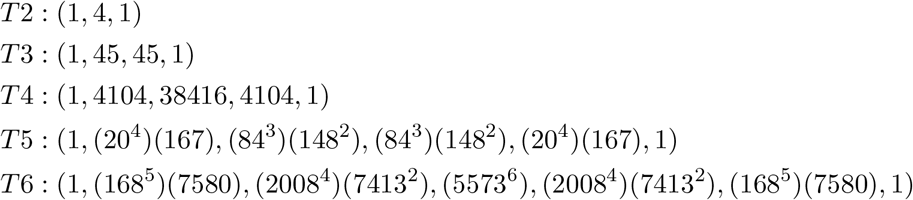

Note that these numbers confirm the observations from the Ising update studies cited above and suggest that the network structure, rather than the choice of the Ising update, is responsible for these results. The methodology is described in detail in the Methods section.

These systems are also multi-stable in nature; perturbing a steady state by flipping one node from ON to OFF or vice versa can lead to a switch to one of the other possible steady states. For even values of n in Toggle-n networks, the majority of the perturbations go back to their original state; for odd value of n, however, the perturbation can lead them to adopt a state at a hamming distance of 1 (Figure S1).

Introducing self-activation and self-inhibition mechanisms into the networks had notable effects on steady-state patterns. The pattern remains unchanged upon the addition of self-activation for three (T3SA), five- (T5SA), and seven-node (T7SA) networks. However, a distinct pattern is seen for even values of n. For the two-node network (T2SA - a toggle switch with self-activation), while 1-high states continue to be most dominant, 0-high (both nodes are OFF) and 2-high (both nodes are ON) also appear. Similarly, for four- (T4SA), six- (T6SA), and eight-node (T8SA) networks, the combinations of 1-high and 3-high, 2-high and 4-high, and those of 3-high and 5-high appear too (Figure S2). Conversely, upon the addition of self-inhibition, the pattern remains the same for two (T2SI), four (T4SI), six (T6SI), and eight-node (T8SI) networks. However, for the three- (T3SI), five- (T5SI), and seven-node (T7SI) networks here, no steady states were observed (Figure S3). When examining the state transitions graphs in these networks, we observe that the states that were stable (i.e., there were no outgoing edges to other nodes) in the cases without self-regulation are able to transition to other previously observed stable states while maintaining their self-edges. We will call such states “meta-stable” (Figures S4A and S4B). The new transitions occur because the self-inhibition on the nodes that are low/off facilitates them to transition to states where they are high. However, in the case of even networks with self-inhibition, the inhibition from the nodes that are high could not be overcome by this mechanism, rendering these states stable (Figure S4C).

Next, we have looked at how the frequency of states with k-high nodes, *F* (*k*), changes with the number of nodes in the network. Thus, *F* (1) would indicate the terminal phenotypes with one TF active. It follows from Theorem 1 that *F* (1) drops sharply from 1 for two nodes to 0.5 for three nodes and 0 for four nodes and more, i.e., single-positive (1-high) states are not observed for networks beyond the toggle triad. The addition of self-activation was able to rescue *F* (1) only for the four-node network (Figure 2A). The direct relevance of Theorem 1 is evident when we consider 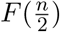 for networks with an even number of nodes; the value is exactly 1 for the networks with no self-regulation and for networks with self-inhibition. The networks with self-activation lose some 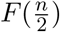 to the states with Hamming distance 1, i.e., the frequency of “flanking states”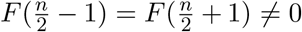. Interestingly, the frequency of these “flanking states” seem to increase with increasing number of nodes (Figure 2A). Similarly, it follows from Theorem 1 that the networks with an odd number of nodes have equal values of 0.5 for 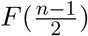 and 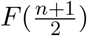.

**Figure 2:**
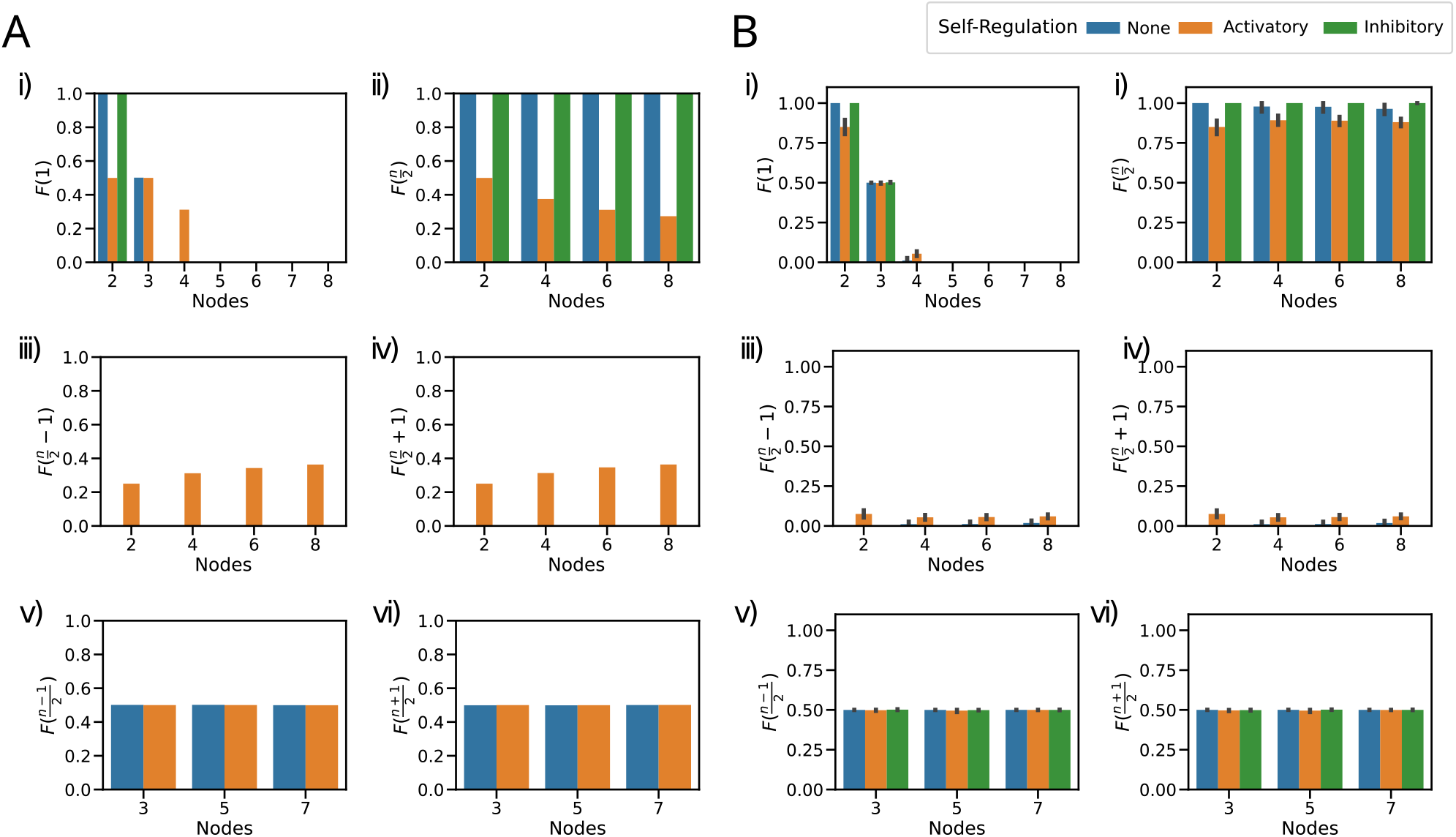
Steady state distributions are dominated by states with n/2 nodes high: A: Frequency of k-high state (*F* (*k*)) for (i) *F* (1) single node high, (ii) *F* (*n/*2) nodes high for even networks, (iii) *F* (*n/*2 − 1) nodes high for even networks, (iv) *F* (*n/*2 + 1) nodes high for even networks, (v) *F* ((*n* − 1)*/*2) nodes high for odd networks and (vi) *F* ((*n* + 1)*/*2) for odd networks. B: Frequency of k-high state (*F* (*k*)) for toggle-n networks with random edge-weights ∈ *U* (0, 1). (i) *F* (1) single node high, (ii) *F* (*n/*2) nodes high for even networks, (iii) *F* (*n/*2 − 1) nodes high for even networks, (iv) *F* (*n/*2 + 1) nodes high for even networks, (v) *F* ((*n* − 1)*/*2) nodes high for odd networks and (vi) *F* ((*n* + 1)*/*2) for odd networks. The error bars represent the 95% confidence interval.

Overall, the 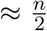 high steady states are predominant for a *n* node mutually repressive regulatory networks instead of single-positive ones that denote (terminally) differentiated states. This could indicate that such networks could govern the differentiation into precursor lineages or progenitor phenotypes with more than three terminal phenotypes. Other regulatory mechanisms would then, therefore, be required for the cell to fully differentiate into terminal phenotypes.

### 2.2 Steady state patterns are robust to edge perturbations

Our analysis so far implicitly provides equal weighting to each edge in the network. To assess the resilience of observed steady-state patterns against extrinsic biological noise (Elowitz et al., 2002), we introduced some variations to the edge weights, reflecting heterogeneity in the strength of regulatory interactions (Figure S5A). Specifically, we sampled the magnitude of each regulatory link from a uniform distribution *U* (0, 1), deviating from the fixed value of 1. We generated 100 such unique sets of edge weight values, each combination denoting a different set of values for varied strength of regulatory interactions among different nodes in that network.

Interestingly, despite these perturbations, the overall pattern of steady states remained largely unchanged, with only minor variations observed. For instance, while the frequency of *F* (1) showed only minor differences compared to the equal-weighted case, notable differences included T4 and T3SI. The pattern becomes more clear when looking at 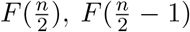 and 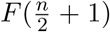. The states with 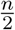 high are more frequently observed in random-weighted networks compared to equal-weighted networks of the same size with self-activation. In contrast, minor variations could occasionally push the 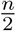 states to the states with Hamming distance =1 (“flanking” states) for the networks without self-regulation. Similarly, the patterns observed in 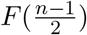 and 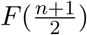 remained consistent, albeit with slight variations, in networks without self-regulation as well as in networks with self-activation (Figure 2B). Interestingly, in circuits with self-inhibition, we observed that the system converges to specific steady states, unlike the case with equal-weighted edges. In certain scenarios, the random sampling chooses weights that do not allow for transition between some of the “metastable” states and thus stabilizing them (Figures S4B, S4D and S4E).

We then explored if perturbations of the sign of regulation would have any effect on the steady-state distributions (Figure 3A). For a low extent of “impurity”, i.e. the replacement of an inhibition by an excitation, the *F* (2) and *F* (3) corresponding to 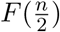 for 4-node and 6-node networks maintain their frequency of 1. Likewise, the *F* (1) - *F* (2), and *F* (2) - *F* (3) corresponding to 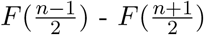 for 3- node and 5-node networks maintain their frequency of 0.5 each for low impurity (Figure 3B), suggesting that the system dynamics exhibits functional resilience to minor changes in network topological space.

**Figure 3:**
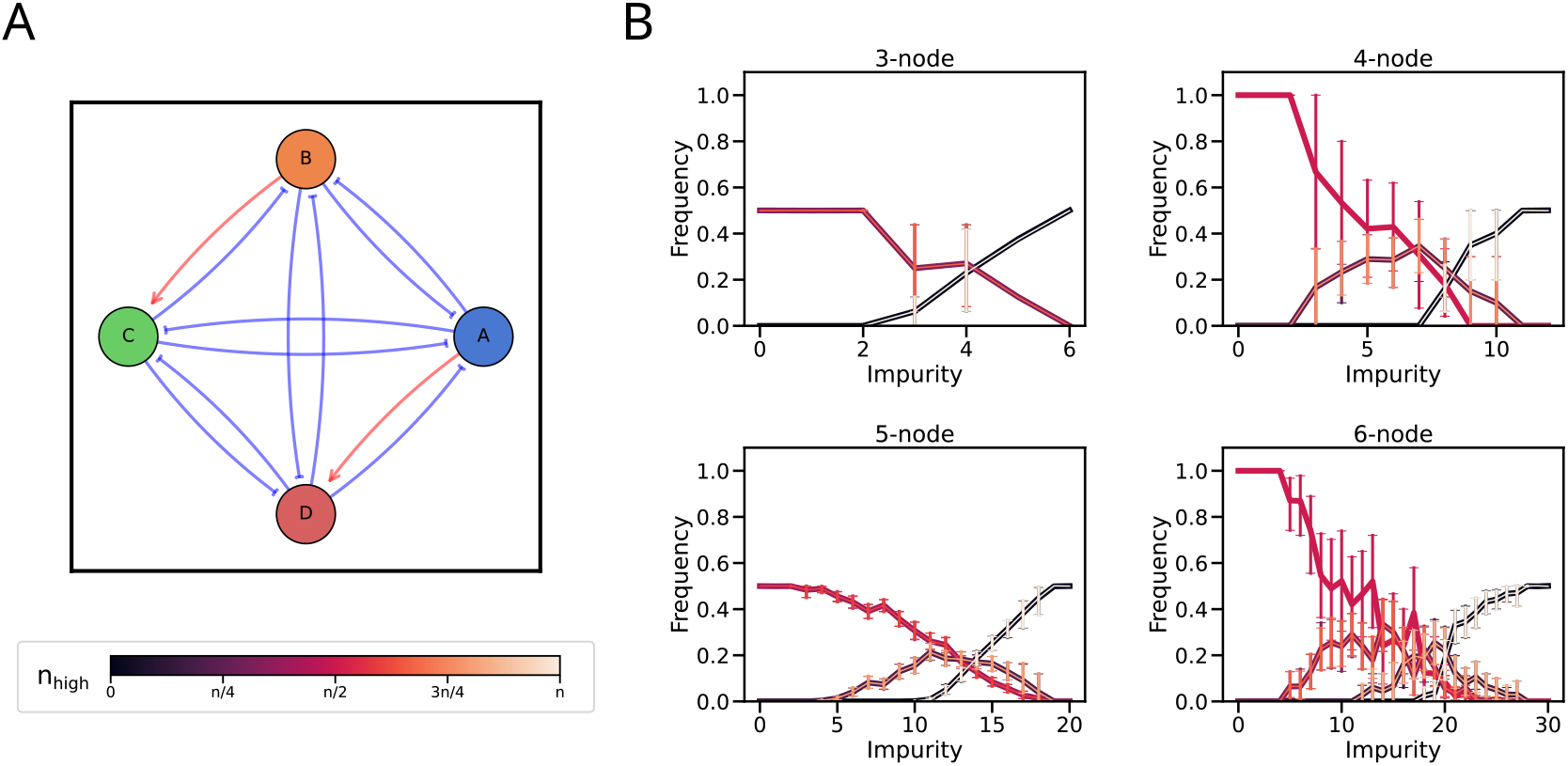
Edge-sign perturbations maintain the steady state distributions for low levels of impurity: A: Schematic of edge-sign perturbation for a 4-node network with 2 impurities. B: Frequency of k-high state (*F* (*k*)) for (i) 3, (ii) 4, (iii) 5, and (iv) 6-node networks where each inhibition is replaced by activation (Impurity). The error bars represent the 95% confidence interval.

As the extent of impurity increased, the steady-state frequency patterns began to change. For even values of n, the frequency distribution shifts to 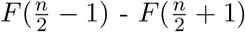 and then broadens to 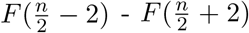, while for odd values of n, it shifts to 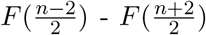. Finally, as expected, *F* (0) and *F* (*n*) become the most dominant states with a frequency of 0.5 each at the maximum level of impurity (i.e. where all inhibitions have been replaced with activation). However, *F* (1) does not exceed 0.5, irrespective of the extent of impurity.

In summary, despite perturbations in edge weights and signs of regulation, steady-state patterns of toggle-n networks exhibit robustness, with only minor variations observed, suggesting resilience to intrinsic biological noise.

### 2.3 Mutually repressive regulatory networks maintain their behavior when embedded in random networks

In biological systems, genes seldom operate in isolation but rather form complex interaction networks, and network motifs such as toggle switch are embedded in larger networks with multiple inputs converging to these motifs. To understand how toggle-n networks behave within such intricate biological contexts, we embedded them (see Methods) into 100 random networks of varying embedding size (i.e, number of nodes in the random network) and embedding density (i.e, average number of edges per node in the random network) (Figure S5B).

We observed that the fundamental behaviors of toggle-n networks remain largely unchanged when integrated into these random networks. We observed a consistent pattern where the frequency of *F* (1) decreased as the number of nodes in the mutually repressive regulatory network increased (Figure 4A), similar to the observations for these networks under isolated conditions. Notably, T4, T5, and T6 deviated from this pattern by displaying non-zero *F* (1) values, though relatively low. The frequency of 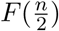 remained the highest for even networks, followed by the states at a hamming distance of 1 (Figures 4B to 4D). Similarly, for odd networks, 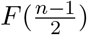 and 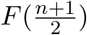 maintained the highest frequency (Figures 4E and 4F). Interestingly, we noted that the frequency of these dominant states 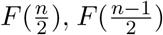, and 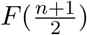 reduced with increasing embedding size and, to a lesser extent, with increasing embedding density.

**Figure 4:**
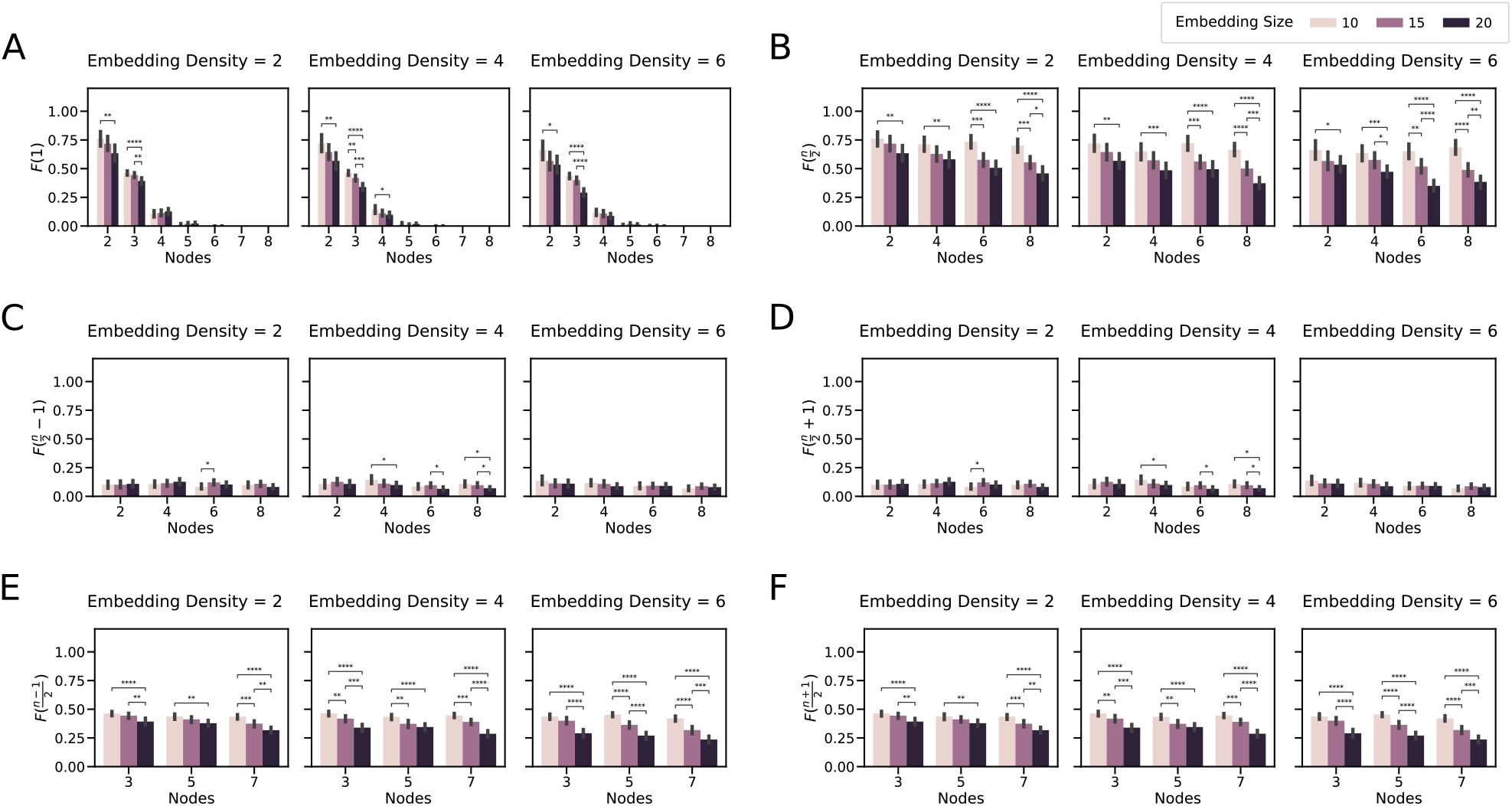
Steady state distributions of the toggle networks are preserved when embedded in random networks: Frequency of k-high state (F(k)) for networks embedded in random networks. The colors represent embedding size (Number of nodes in the random network), and the columns represent embedding density (average number of edges in the random network). A: F(1) single node high, B: F(n/2) nodes high for even networks, C: F(n/2-1) nodes high for even networks, D: F(n/2+1) nodes high for even networks, E: F((n-1)/2) nodes high for odd networks and F: F((n+1)/2). The error bars represent the 95% confidence interval.

Despite the random interaction among networks, the overall outcomes of toggle-n networks remain consistent with that observed in isolation. It is also worth noting that we did not control for any specific network architecture. For example, a network interacting in a hierarchical manner with the toggle-n networks and could potentially influence their behavior in a directed manner (Krumsiek et al., 2011; Peter & Davidson, 2017).

### 2.4 Network of mutually inhibiting teams of TFs mirror the toggle networks

To check if these simplified toggle-n networks represent the behavior of larger networks where a group of TFs regulates a particular phenotype, we considered “Team-n” networks. These are fully connected networks with n teams such that the nodes within a team have activatory edges and nodes across teams have inhibitory edges (Figure S5C). A team *T*_*i*_ is said to be high when the expression of all the members in a team is high.

As a start, we assumed the number of members (*m*) to be equal between the teams. The *F* (1) matches the exact same frequency of the toggle-n networks regardless of the number of members (Figure 5A). Comparing the 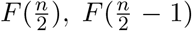 and 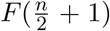 for the networks with even number of teams with those of toggle-n networks, we see that the pattern matches with the networks with either no regulation or self-inhibition (Figures 5B to 5D). Similarly, comparing 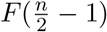 and 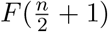 for odd networks narrows it down to be equivalent to the toggle-n case without self-regulation (Figures 5E and 5F).

**Figure 5:**
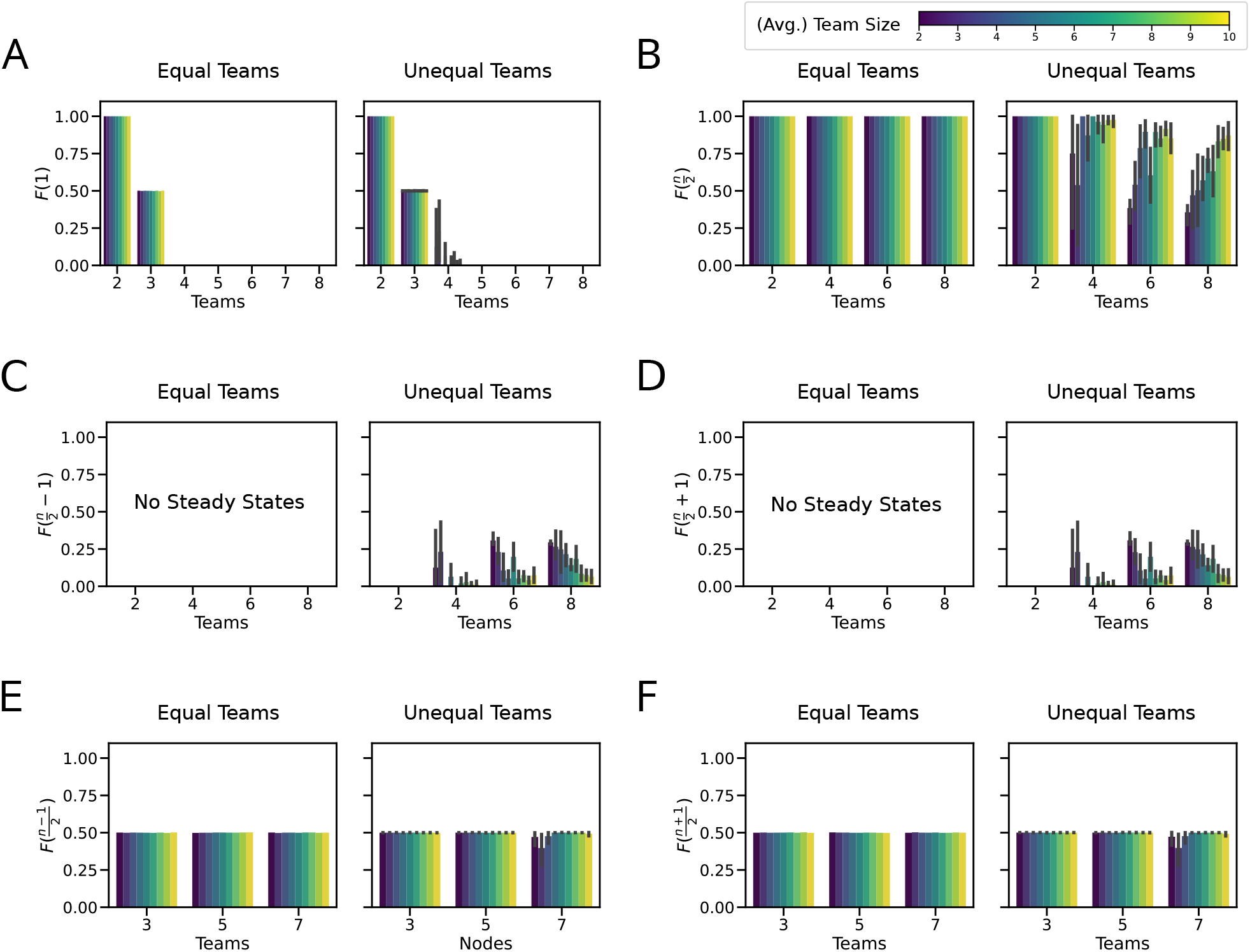
Steady state distributions of mutually repressing team network mirror the toggle networks: Frequency of k-high state (F(k)) for Team-n network of varying team sizes. Left column: Team sizes are equal for all teams, Right column: Team sizes are different across teams. The colors represent the average team size. A: F(1) single node high, B: F(n/2) nodes high for even networks, C: F(n/2-1) nodes high for even networks, D: F(n/2+1) nodes high for even networks, E: F((n-1)/2) nodes high for odd networks and F: F((n+1)/2). The error bars represent the 95% confidence interval.

However, there can be cases where the number of members in a particular team can vary across teams (Chauhan et al., 2021). So, we have also considered cases by randomly splitting the members between the teams. The *F* (1) in this case, while dropping with an increasing number of teams, also shows a non-zero frequency with the four-team network compared to the equal teams case (Figure 5A). However, this frequency shows a high variance between the networks considered, and the frequency also drops with increasing average team size *m*. Similarly, while the behavior for even networks remained largely unchanged, with the 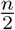 states still being the dominant states, the states with a hamming distance of 1 from it 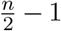 and 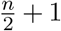 seem to steal some of the frequency away from it for low average team sizes, again with a high variance (Figures 5B to 5D). The 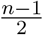 and 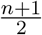 states also remain similar to the team-n case for networks with an odd number of teams(Figures 5E and 5F). This is reminiscent of the perturbation of edge weights, and the fraction of members in a particular team could be considered equivalent to the relative strength of interactions between the teams. The high variance for networks with a low average team size could be attributed to the larger differences in team sizes, which create more discrete levels. As the average team size increases, we expect these *F* (*k*) values to converge to the levels seen with edge-weight perturbations, because the differences in team size become smaller, resulting in more continuous levels.

These results show that toggle-n networks can be used to represent larger networks of TFs interacting in a team-like manner.

### 2.5 Multilevel formalism reveals hybrid states characterized by intermediate expression levels

After thoroughly examining the phenotypic spaces of “Team-n” networks, we asked if expanding the state space of the simulation formalism would unveil any additional characteristics of these networks. We simulated fully connected Team-n networks with equal number of members in each team, using a set of multi-level models defined by Equation (4). Briefly, the multi-level formalism expands the state space by increasing the number of expression levels each node can have. For example, in the four-level model, node expressions take four values – {-1, -0.5, 0.5, 1} instead of two {-1,1} in the traditional Boolean formalism. To characterize the steady states of the multi-level model, we used team scores (average expression of all nodes in a given team). As the nodes in each team showed identical expression levels, the team score perfectly reflects the expression of every node in the team (Figure S6).

Recently, we found that simulating GRNs underlying Epithelial-Mesenchymal Plasticity (EMP) using four-levels uncovers new hybrid states (states where nodes from both E and M teams have non-zero expression) in the phenotypic landscape (Hari et al., 2024). However, simulations of the two-team network using a four-level model revealed no new steady states (Figure 6A). While EMP networks are also two-team networks, some nodes do not completely obey the team structure, causing “frustration” in the network structure, a property absent in the two-team network simulated here. A network is structurally frustrated if there exists at least one pair of nodes in the network such that not all the direct and indirect interactions in the network are of the same sign. Note the in two-team network, every pair of nodes belonging to the same team is connected by only positive direct and indirect (mediated by other nodes in the network) interactions and vice-versa. Therefore, the two-team network is not frustrated. Because such frustration can lead to an increase in the fraction of hybrid states (Rashid et al., 2022; Tripathi et al., 2020), we attribute the lack of new hybrid states in the two team networks to the lack of frustration. While a two-team network is not frustrated, every Team-n network with *n >* 2 is frustrated. Consider three nodes A, B, and C in a three-team network, each belonging to a different team. Since the networks are fully connected, these nodes mutually inhibit each other. Considering the interactions between A and B, the direct interactions both have a negative sign, but the indirect interaction mediated by C consists of two connected inhibitions resulting in a net positive interaction. This reasoning holds true for any pair of nodes in Team-n networks (*n >* 2). Thus, we expect to see new hybrid states when Team-n networks are simulated using the multi-level formalism.

**Figure 6:**
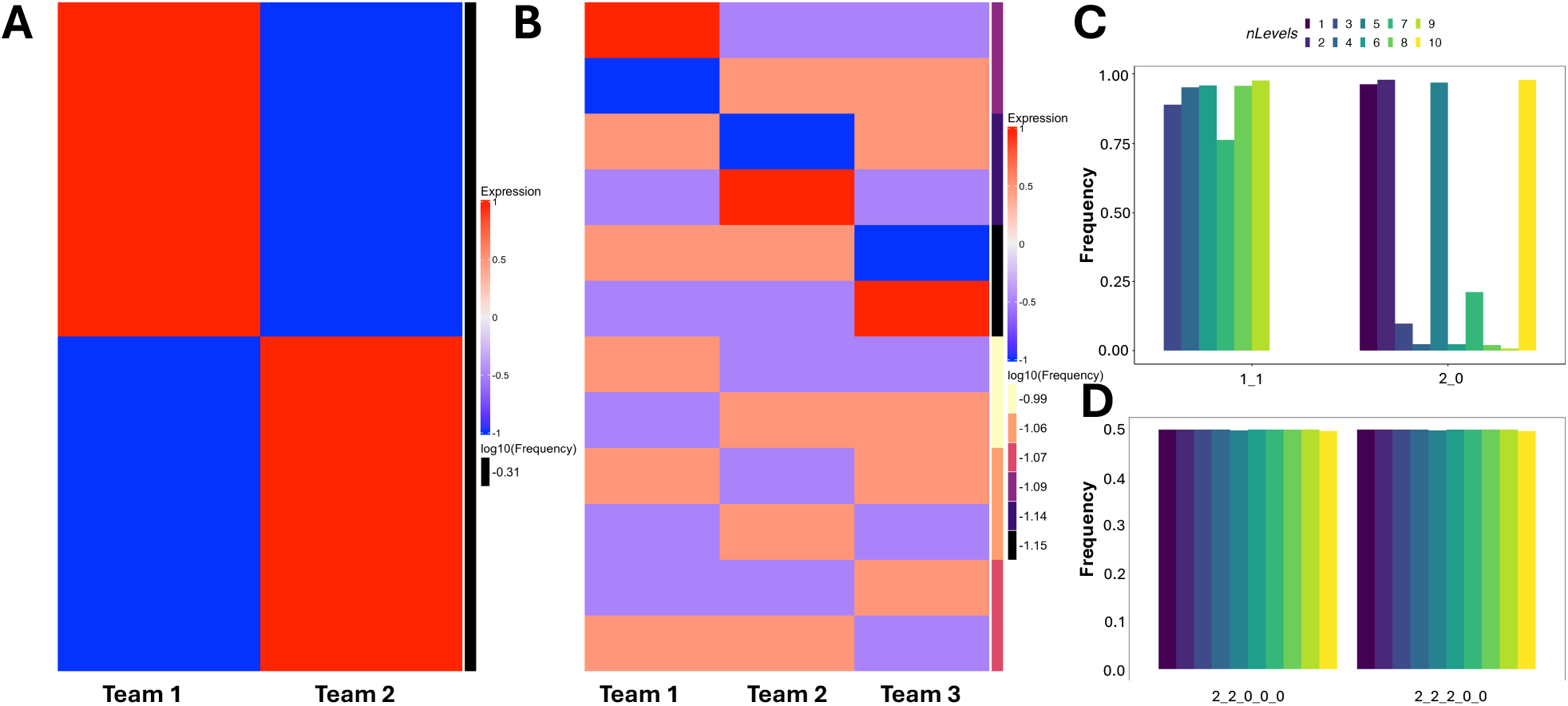
Steady states resultant from multilevel formalism mirror the toggle networks but with partial team scores: A: Team scores for the steady states of 2 team network simulated using four-level model. B: Same as A, but for 3 team network. C: normalized and discretized steady state configurations for two team network for different levels of the multilevel formalism. D: Same as C but for 5-team network.

In contrast to a two-team network, a three-team network results in an entirely new set of steady states when simulated with multilevel models. While the configuration of states remained the same (high-low-low and high-high-low Figure 6B), all steady states of the three-team network show partial expression in at least two out of three teams. As the number of levels increases, all teams show partial expression in all steady states. The maximum expression level of each node also decreases as the number of levels increases (Figure S7A). This decrease in expression level was observed across networks with four to ten teams (Figure S7).

We then asked if the partial expression of teams is purely because of the decrease in the maximum possible node expression. We normalized the expression of each node by dividing the expression level by the maximum possible expression in the simulation and categorized the team scores into 0, 2, or 1, corresponding to a normalized expression of -1, 1, or between -1 and 1, respectively. For two and three-team networks, we found that most steady states have partial team expression, even after the normalization, such that the steady-state configuration with the highest frequency was 11 for the two-team network and 111 for the three-team network (Figures 6C and S8A). We found similar dominance of partial expression in the four-team network as well (Figure S8B). However, from the five-team onwards, we find that even with more levels in the allowed state space, all team scores were either 2 or 0 (Figures 6D and S8C to S8F). More than 99% of the state space converged to the steady states expected from the team configurations (22000 and 22200 for five-team networks, 222000 for six-team networks, and so on).

### 2.6 Toggle-n networks require the synergy of epigenetic reprogramming and cytokine signaling to adopt terminal fate

To further investigate the mechanisms that cells can adopt for making robust cell fate choices, we introduced asymmetry in the strength of transcriptional links in the networks. This asymmetry could arise from variations in DNA methylation, histone acetylation, or mRNA degradation (Duddu et al., 2022). We hypothesized that stronger inhibition of TF A on the other cell state-specific TFs would direct cells toward a cell state characterized by high expression of transcription factor A. Therefore, we focused on the frequency of the single positive state where only transcription factor A is highly expressed, *F*_*A*_(1).

We increased the inhibitory link strengths from A to all remaining nodes (referred to edge weights hereafter) from 1 to 10, while keeping the rest at a default value of 1, and analyzed the steady-state distributions of cell states. We expected that higher edge weights would be able to have *F*_*A*_(1) = 1. However, we observe that the steady state distribution of *F*_*A*_(1) saturates at 0.5, with the remaining states exhibiting co-expression of other cell-state specific TFs along with A (Figure 7A).

**Figure 7:**
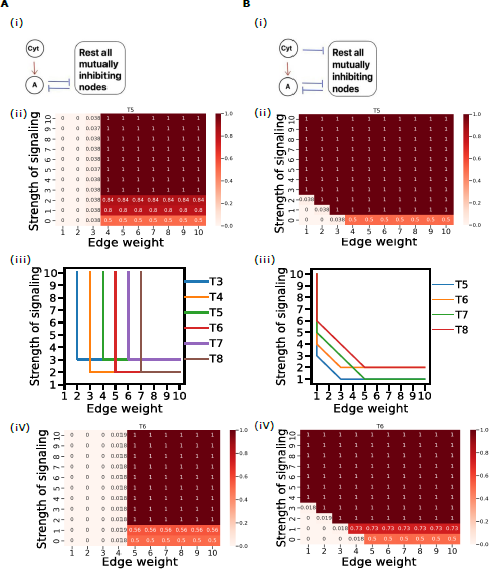
Relative frequency of single positive A-high, *F*_*A*_(1) with increasing strength of signaling and edge weight: A: (i) Network diagram representing cytokine (Cyt) activating the TF A. (ii) Heatmap representation of the relative frequency of the cell state *F*_*A*_(1) with increasing edge weight and strength of signaling for a representative case of toggle pentagon. (iii) Line plot showing the threshold values of the strengths of signaling for a range of values of edge weights and the threshold values of edge weights for a range of values of strengths of signaling for observing *F*_*A*_(1) with a relative frequency of 1. (iv) Heatmap representation of the relative frequency of the cell state *F*_*A*_(1) with increasing edge weight and strength of signaling for a representative case of toggle hexagon. B: Same as A but for the scenario in which the cytokine inhibits other TFs in addition to activating TF A.

Next, we investigated the role of cytokine signaling in cellular differentiation. We hypothesized that a cytokine that transcriptionally activates A would push the cells toward the single positive A high state. We introduced a cytokine that constitutively activates A in our simulations and varied the strength of signaling (strength of activation of A by the cytokine) from 0 to 10 and measured *F*_*A*_(1) as before. Contrary to our expectations, we observed no significant changes in the distribution of the cell state with increasing signaling strength (Figure 7A).

We then combined increased edge weights (corresponding to epigenetic reprogramming) with varying strengths of cytokine signaling and ran the simulations again. The results suggest that a threshold level of both signaling strength and edge weight is required for complete differentiation towards a single positive A-high state. This means that both cytokine signaling and epigenetic reprogramming are necessary simultaneously, and one cannot compensate for the absence or deficiency of the other. The threshold for edge weight increases linearly with the size of the network. For a network with three nodes (T3), the edge weight threshold is 2; for T4, it is 3; for T5, it is 4; for T6, it is 5; for T7, it is 6; and finally T8, where the threshold is 7. In contrast, the threshold for signaling strength only depends on whether the network has an odd or even number of nodes. For networks with an odd number of nodes (T3, T5, T7), the signaling strength threshold is 3, whereas, for the networks with an even number of nodes (T4, T6, T8), the threshold is 4 (Figure 7A).

Finally, we have included inhibitory edges from the cytokine to other nodes, keeping the existing activatory link on A to see if this scenario would be able to give only single positive A high state. Consistent with our hypothesis, we observed only the A-high state (*F*_*A*_(1) = 1) in the steady state distribution with a sufficient increase solely in the strength of signaling (both strength of activation on A by the cytokine and strength of inhibition on the other nodes by the cytokine). The signaling strength required for *F*_*A*_(1) = 1 in this case increases linearly with network size. For networks with five nodes (T5), the signaling strength threshold is 3; for T6, it is 4; for T7, it is 5; and for T8, the threshold is 6 (Figure 7B). Similarly in this scenario, increased edge weight can compensate for the deficiency of signaling strength and the signaling strength threshold for *F*_*A*_(1) = 1 decreases with increasing edge weight (Figure 7B).

## 3 Discussion

Through our Boolean modeling approach, we aimed to unravel the dynamics of toggle-n networks and their ability to replicate the intricate process of cellular differentiation observed in biological systems. However, our findings suggest that these networks may not efficiently generate terminally differentiated states, through the frequency of single positive states *F* (1) or lack thereof. The predominance of “central states”, indicated by ≈ *n/*2 for even networks and (*n* − 1)*/*2 - (*n* + 1)*/*2 for odd networks, suggests a bifurcating process rather than a direct path to terminal differentiation.

This bifurcating process may be more reflective of initial differentiation to precursor lineages or progenitor cells which further differentiate to the terminal phenotypes (Valencia & Peter, 2024). For instance, in the case of haematopoietic stem cells, these can differentiate into progenitor cells committed to specific lineages, such as myeloid precursor and lymphoid precursor lineages. These lineages can further differentiate into erythrocytes, macrophages, neutrophils, basophils, eosinophils and platelets, and into B cells, T cells, and natural killer cells, respectively (Laurenti & Göttgens, 2018; Zhou & Huang, 2011).

Next we went on to check on what sort of conditions these networks would be resilient to, and the conditions that can aid towards the terminal differentiation. We observe that when subjected to random noise through perturbation of the edges both in-terms of their magnitude as well as sign, the networks maintained their traits. Similarly, the traits are also maintained when such networks interact with external signals from random networks. However, this behavior changes when these mechanisms act in an asymmetric or biased manner. The presence of selective activatory external signal to one particular node (in this case A) or increased strength of inhibition from that node to other nodes are able to shift the stable states towards a single positive state involving that particular node *F*_*A*_(1). Additionally, these mechanisms act in a synergistic manner and contribute towards a higher *F*_*A*_(1) when acting in unison. These simple toggle-n GRNs also remain scalable when the TFs interact in a teams-like manner as observed in various networks of cellular differentiation.

It is plausible that the actual gene regulatory network (GRN) governing differentiation may exhibit a structure vastly different from what we have explored. Such GRNs can be inferred and constructed from experimental developmental data (Kim et al., 2024; Okawa et al., 2015, 2016). These GRNs could provide a more nuanced and biologically relevant understanding of the developmental process. However, our study has focused on a simplified scenario, where each cell state-specific TF inhibits every other, thereby scaling up the well-established motif of a toggle switch. Moreover, there is a possibility that the topology and strength of regulation within the GRN could be dynamic traits themselves. While we have investigated discrete cases of remodeling regulatory strengths, it is conceivable that these regulations are subject to dynamic alterations and reinforcement through feedback mechanisms from specific nodes. Nevertheless, such a comprehensive modeling effort falls beyond the scope of our current study.

Furthermore, the choice of a Boolean framework over a more continuous representation of states introduces its own set of implications. This choice was, however, justified due to the the problems of defining a high state and discretizing the steady-state distributions for the context of our simulations. Even after relaxing the Boolean framework to allow more granular expression levels, we observe that the trends remain the same. But it is worth noting that the undifferentiated stem-like state is not well defined. Even if we consider an all-high or all-low state to be representative of a stem-cell it is not a stable steady state. Moreover, the differentiation process is also a continuous process involving gradual transitions influenced by varying levels of transcription factors. Continuous frameworks such as ODEs are more suited to capture such dynamics (Lu et al., 2025).

The emergence of n/2-high stable states is a consequence of both the structure of the repressive networks we study and the Ising update function. We prove that the update dynamics of such a function always converge to one of the n/2-high stable states. It is plausible that alternative formalisms, such as those employing logical AND/OR gates, could yield vastly different results (Andrecut, 2011; Gedeon, 2024b). The selection of an appropriate Boolean function to represent regulations within mutually repressive networks remains a subject of debate. To address a concern about the generalizability of the reported results, we have also considered all possible monotone boolean functions (MBFs) compatible with signs of network edges and calculated the number of those functions that support a steady state with k-high states 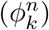. Our calculations show that among all monotone boolean functions the majority support the n/2 high steady state. However, this analysis cannot be done for larger networks as the number of MBFs (Dedekind number) grows extremely quickly. However, for large networks we can always analyze samples from the collection of all MBFs. On the other hand we can replace Ising update model by selecting other monotone Boolean functions, perhaps those which have high likelihood of occurence in biological systems (Kadelka & Murrugarra, 2024; Subbaroyan et al., 2022). Our work provides valuable insight into the mechanistic aspects of directed differentiation of stem cells. Such studies would have profound implications in fields like synthetic biology and regenerative medicine. By inducing these networks in prokaryotic or eukaryotic cells, depending on the context and application, a desired type of cells can be synthesized for therapeutic purposes, for example, in the synthesis of probiotics for treating certain kinds of bowel diseases and in engineering T cells for use in cancer immune therapy. Understanding mechanistic aspects of cellular differentiation would also have profound implications in regenerative medicine.

## 4 Methods

### 4.1 Boolean Formalism

We simulated the dynamics of the networks using asynchronous Boolean simulations. We used the Ising formalism, a well-established framework in physics (Bortz et al., 1975) and which has been recently applied to the study of regulatory networks (Font-Clos et al., 2018; Hari et al., 2022a).

For a network with n-nodes, the state **x** ∈ {−1, +1}^*n*^, is a n-dimensional vector. Here, *x*_*i*_ = +1 indicates the gene *i* to be in a ON/high state and *x*_*i*_ = −1 indicates the gene *i* to be in a OFF/low state. The choice of -1 over 0 for the low state is to allow a particular gene to contribute towards the regulation of the target nodes even in the low state.

The topology files which are the edges listed in tab-separated format were used as input for the simulation. During simulation these were converted and represented as adjacency matrices **Adj**. The elements of **Adj** are such that:

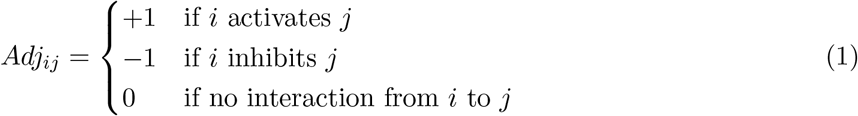

These simulations were performed by generating 10000 initial conditions for each network topology. The updating was done using asynchronous update where *k* is chosen randomly in {1, 2, … *n*} and the *k*^th^ node is updated. This update was done for a maximum of 1000 time steps or until convergence to steady state. The update rules are as follows:

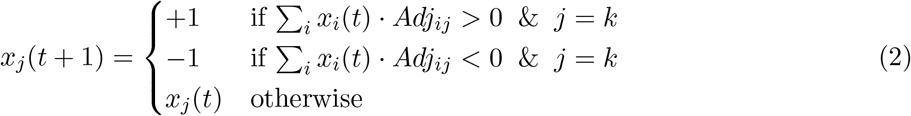

The simulation outputs the steady states and their average relative frequency *f* (*i*) across three replicates. The steady states that do not converge at the end of maximum time steps are not considered further. The frequency of *k* − *high* states, *F* (*k*) is given by,

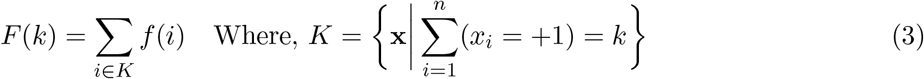

We also simulated the Team-n networks with multi-level extension of the ising formalism, governed by the following set of update rules:

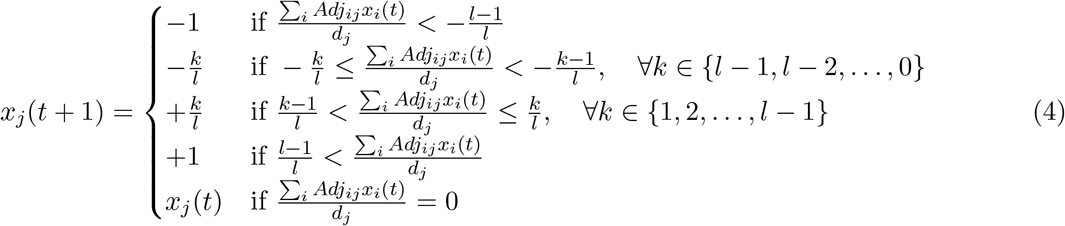

Where *l* is the number of levels defining the state space’s granularity and *d*_*j*_ is the in-degree of *j*^*th*^ node.

### 4.2 Network Generation

#### 4.2.1 Toggle-n networks

We generated networks using the NetworkX library (Hagberg et al., 2008) to create a series of togglen networks with varying numbers of nodes. Toggle-n networks are mutually repressive regulatory network, where each node inhibits the other nodes. To generate these networks, we created complete directed graphs for nodes ranging from 2 to 8 giving T2 to T8. The edge weight parameter for all edges was set to 2, representing inhibition. For networks with self-regulations, self-edges are added to the graph with edge weight parameter = 1 or edge weight = 2 for self-activation or self-inhibition cases, respectively. These were converted to edge-list format and saved as a tab-separated topology file (.topo file).

The adjacency matrix **Adj**(*Toggle*) in this case would be:

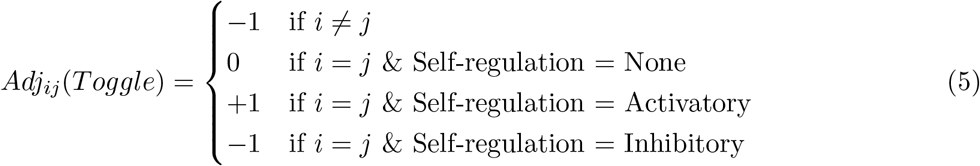

#### 4.2.2 Addition of impurities

We started with a particular Toggle-n network and replaced *n*_*imp*_ possible combination of inhibitory edges with activations for *n*_*imp*_ ∈ 0, 1, … *n*(*n* − 1). However, since most of the networks end up being equivalent due to symmetry, we checked if any of these graphs are isomorphic and selected a maximum of 100 such non-isomorphic graphs for *n* ≤ 5 and 50 for *n* = 6. For *n >* 6, the number of combinations become too high and this analysis was not performed. The isomorphism testing was performed in a manner similar to that in the previous work (Duddu et al., 2024). For this, the complete directed-weighted graph was reduced into an equivalent directed-unweighted graph with the presence of an edge representing an activation and the lack representing an inhibition. The isomorphism was checked using nx.algorithms.is isomorphic function which utilizes an implementation of the VF2++ algorithm (Foggia et al., 2001). These were then saved as topology files.

#### 4.2.3 Embedding toggle networks in random networks

We embedded toggle-n networks in random directed graph networks. For each combinations of embedding sizes (the number of nodes in the random network) of 10, 15 and 20, and embedding density (the average number of edges to each node) of 2, 4 and 6, 100 random directed graphs were generated using nx.gnm random graph. The edge-weights for edges present in this network was randomly sampled from {1, 2}. This network was merged with each of the toggle network considered. To connect the two networks together, all the edges between nodes of the random network and toggle network are randomly sampled from {0, 1, 2}.

#### 4.2.4 Team-n networks

A Teams-n network consists of a directed graph with n-teams with nodes in each team (*T*) having intra-team activations and inter-team inhibitions. This is done by having an adjacency matrix **Adj**(*Team*), such that:

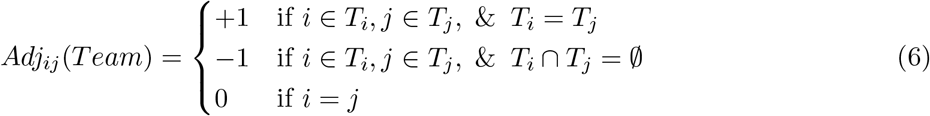

We have considered the cases of both equal members in each team and random unequal split of members between the teams. For the equal case, the number of members in a team, 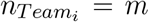, where *m* ∈ {2, 3, … 10}. For the unequal case, 100 network were generated where the *m*_*i*_ is sampled from Multinomial distribution with 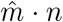 number of experiments and *p*_*i*_ = 1*/n*, while ensuring that *m*_*i*_ ≠ 0 for any *i* and 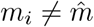 for all *i*.

Since, the steady states only allow all members of a team to be either all high or all low, the results would be consistent even with a weaker condition, such as average expression being greater than 0.5.

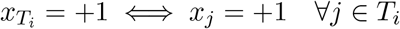

### 4.3 Edge weight perturbation

To simulate the effect of biological noise acting through the strength of regulatory interactions, we considered each edge to have random weights. For each topology, we performed 100 sets of edge weight sampling, where each element of the adjacency matrix was scaled by a randomly sampled variable from a uniform distribution between 0 and 1, *U* (0, 1). This can be represented by the adjacency matrix **Adj**(*E*.*W*.):

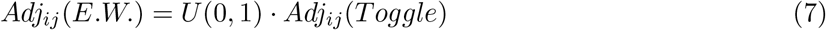

Further simulations were then performed as mention in Section 4.1.

Similarly, to assess the role of asymmetry of regulation strength in directed differentiation towards terminal fates, we altered the strengths *W* of inhibitory edges (referred to as edge weight hereafter) from node A to all remaining nodes when that node is high. In other words, if A is inhibiting B and has a edge weight of *W >* 1, then node A would contribute −*W*, if A is high or +1, if A is expressed low to the net effect of transcriptional regulation of B. This weight *W* was varied from {1, 2, …, 10}. The adjacency matrix for such a case where A is the *k* ^*th*^ node is:

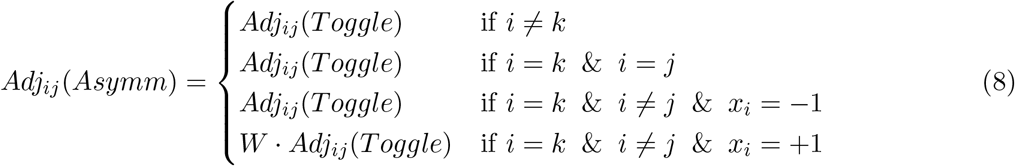

The number of states with only A high, *F*_*A*_(1), was then quantified to assess the impact of this asymmetry in directed differentiation.

### 4.4 Signaling

Next, we wanted to decipher the role of synergy between cytokine signaling and asymmetry in the networks for directed differentiation towards *F*_*A*_(1). We have included a cytokine that activates cell state-specific transcription factor A in the networks by making necessary changes in the topology file (refer to Section 4.1) and performed simulations with increasing strength of signalling (similar to increasing *W* in Equation (8)) and edge weights. Suppose if the transcription factor A is the *k*^*th*^ node and the cytokine is in the *l*^*th*^ node, then the adjacency matrix with an edge weight of *W* and a strength of signalling *W*_*C*_ is given by:

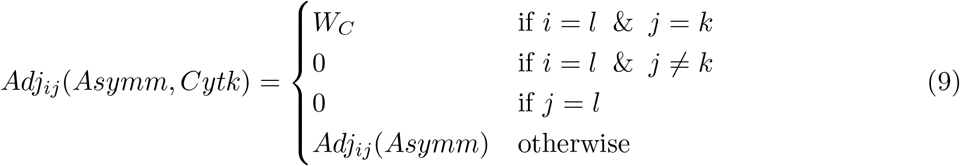

In addition to these, we included inhibitory edges on the rest of the nodes from the cytokine by keeping the existing activation on A intact. Now the adjacency matrix is given by:

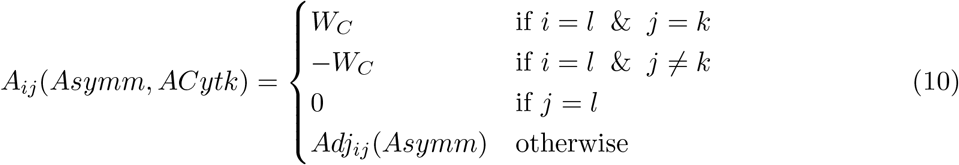

### 4.5 Monotone Boolean functions

**Definition 1**. A function *f*_*i*_ : 𝔹 ^*k*^ → 𝔹 is a *positive (negative) monotone Boolean function* with respect to input *x*_*j*_ if

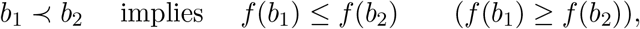

for all pairs *b*_1_ ≺ *b*_2_ ∈ 𝔹^*k*^ which only differ in *j*-th component and 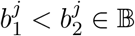.

**Definition 2**. Consider a signed graph (network) of interactions *N* = (*V, E*) with a set of *n* vertices *V* and a set of edges *E*, where each edge *e* = *e*(*i* → *j*) has a sign 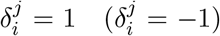 for a *activating (repressing) edge*. A Boolean function *f* = (*f*_1_, …, *f*_*n*_) is *monotone Boolean function (MBF) with respect to network N* (Gedeon, 2024a, 2024b) if each *f*_*i*_ is monotone with respect to all its arguments *x*_*j*_ that respect the sign 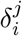, i.e. *f*_*i*_ is positive Boolean function with respect to each *x*_*j*_ where 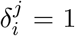 and negative Boolean function with respect to each *x*_*j*_ where 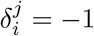.

These definitions highlight that MBFs preserve the order of inputs, ensuring that an increase in input values does not result in a decrease in function’s output.

#### 4.5.1 Formulation of the problem

Consider a series of networks *Tn* with *n* = 2, 3, 4, 5, 6 with *n* nodes and repressive edges connecting all ordered pairs of nodes (*i, j*) ∈ *V* × *V, i* ≠ *j*. Let

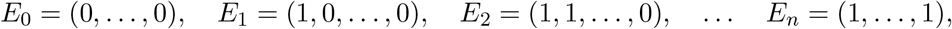

be states in 𝔹^*n*^ with 0, 1, …, *n* ones in the Boolean vector.

Consider a collection ℱ^*n*^ of monotone Boolean functions monotone with respect to network *Tn*. An element of ℱ*n* is a *n*-tuple *f* = (*f*_1_, *f*_2_, … *f*_*n*_) where each *f*_*j*_ : 𝔹^*n*−1^ → 𝔹 is a monotone Boolean function that respects the signs of the network edges. Let 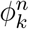 be the number of MBFs *f* ∈ ℱ^*n*^ for which the state *E*_*k*_ ∈ 𝔹^*n*^ is an equilibrium, i.e it satisfies

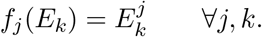

Here 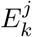 is *j*-th component of vector *E*_*k*_.

Observe that since network *Tn* is invariant under group of permutations *S*_*n*_, the number of functions supporting equilibrum *E*_*j*_ and an equilibrium obtained by any permutation of *E*_*j*_, will be the same.

#### 4.5.2 Approach

**Example** Before we describe how we evaluate numbers 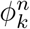, for *n* = 3, 4, 5, 6 and *k* = 0, …, *n* we illustrate our approach on a simple example of toggle switch (Gardner et al., 2000) *T* 2 with nodes 1, 2 and corresponding Boolean variables *x*_1_, *x*_2_. The monotone Boolean functions with respect to *T* 2 are pairs *f* = (*f*_1_, *f*_2_) where both *f*_1_, *f*_2_ : 𝔹 → 𝔹. There are three negative monotone Boolean functions 𝔹 → 𝔹 and thus respect the signs of *T* 2: the zero **0** (one **1**) function, assigning to both inputs the output 0(1) and the −*I* function that assigns 0 to input 1 and 1 to input 0. Therefore ℱ^*n*^ has 3 × 3 = 9 functions *f* = (*f*_1_, *f*_2_). To compute 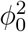, the function *f* = (*f*_1_, *f*_2_) has to satisfy

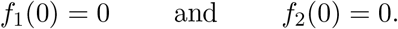

There is exactly one such function *f* = (**0, 0**). Similarly, to compute 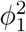 the function *f* = (*f*_1_, *f*_2_) has to satisfy

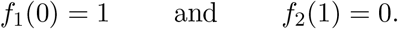

There are two functions {**1**, −*I*} that satisfy the first condition, and two functions {**0**, −*I*} that satisfy the second condition. This gives 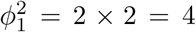. Finally, *f* = (**1, 1**) is the unique function that supports *E*_2_. Therefore

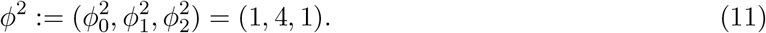

Even in this simple example we observe a “hat”-like pattern where the equilibria with equal number of zeros and ones are supported by more MBFs than equilibria that are more uniform.

We compute 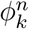, for *n* = 3, 4, 5, 6 and *k* = 0, …, *n* we proceed in three steps, which are implemented in Python.

1. We explicitly construct the collection ℱ^*n*^, *n* = 3, 4, 5, 6 by first constructing all Boolean functions *f*_*i*_ : 𝔹^*n*−1^ → 𝔹, discarding those that do not satisfy negative monotonicity conditions and forming *n*-tuples of such functions *f* = (*f*_1_, …, *f*_*n*_) ∈ ℱ^*n*^.
2. For any input *b* ∈ 𝔹^*n*−1^ we calculate 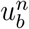 the number of Boolean functions *f*_*i*_ : 𝔹^*n*−1^ → 𝔹 such that

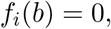

and its complement 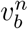, the number of functions *f*_*i*_ : 𝔹^*n*−1^ → 𝔹, such that

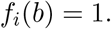

Clearly, 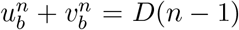 the Dedekind number (Fidytek et al., 2001; Jäkel, 2023) which is the number of all MBFs with *n* − 1 inputs.
3. The previous calculation is simplified by the observation that the MBF *f*_*i*_ : 𝔹 ^*n*−1^ → B is invariant under any permutation of the inputs. Therefore

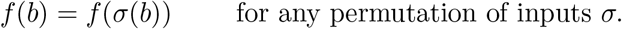

Therefore, the values 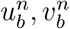 only depend on number of 1s and 0s in the input *b*. We denote these values *u*^*n*^(*j*), *v*^*n*^(*j*) where *j* = 0, …, *n* − 1 is the number of 1s in the input *b* ∈ 𝔹^*n*−1^. These number are calculated by the Python code and are listed in Table 1.

**Table 1:** Values of *u*^*n*^(*j*) and *v*^*n*^(*j*) for *n* = 2, 3, 4, 5, 6.

| Input in $\mathbb{B}^{n-1}$ | $u^n$ | $v^n$ |
| --- | --- | --- |
| $n = 2$ | | |
| (0) | 1 | 2 |
| (1) | 2 | 1 |
| $n = 3$ | | |
| (0,0) | 1 | 5 |
| (1,0) | 3 | 3 |
| (1,1) | 5 | 1 |
| $n = 4$ | | |
| (0,0,0) | 1 | 19 |
| (1,0,0) | 6 | 14 |
| (1,1,0) | 14 | 6 |
| (1,1,1) | 19 | 1 |
| $n = 5$ | | |
| (0,0,0,0) | 1 | 167 |
| (1,0,0,0) | 20 | 148 |
| (1,1,0,0) | 84 | 84 |
| (1,1,1,0) | 148 | 20 |
| (1,1,1,1) | 167 | 1 |
| $n = 6$ | | |
| (0,0,0,0,0) | 1 | 7580 |
| (1,0,0,0,0) | 168 | 7413 |
| (1,1,0,0,0) | 2008 | 5573 |
| (1,1,1,0,0) | 5573 | 2008 |
| (1,1,1,1,0) | 7413 | 168 |
| (1,1,1,1,1) | 7580 | 1 |

In the final step, we directly enumerate 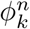 by using numbers *u*^*n*^(*j*) and *v*^*n*^(*j*). We illustrate this for the case *n* = 4. We first compute 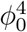. The function *f* = (*f*_1_, *f*_2_, *f*_3_, *f*_4_) supports an equilibrium *E*_0_ = (0, 0, 0, 0) if

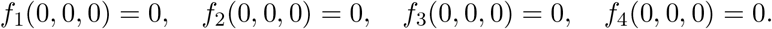

Using the *n* = 4 sub-table in Table 1 we compute the number of functions that satisfy each of these conditions. Since any combination of such functions gives an MBF that supports *E*_0_, we get

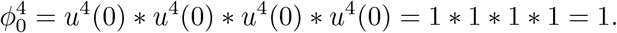

A more interesting is computation of 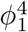. The function *f* = (*f*_1_, *f*_2_, *f*_3_, *f*_4_) supports an equilibrium *E*_1_ = (1, 0, 0, 0) if

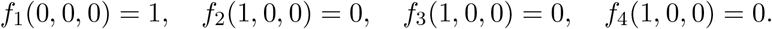

Using the *n* = 4 sub-table in Table 1 we compute

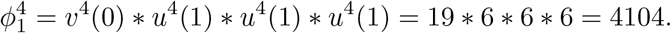

#### 4.5.3 Number of MBF supporting steady states in T3, T4, T5 and T6

##### Network T3

We consider states *E*_0_ = (0, 0, 0), *E*_1_ = (1, 0, 0), *E*_2_ = (1, 1, 0), *E*_3_ = (1, 1, 1). Then the numbers

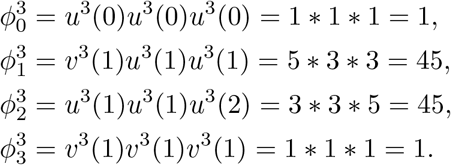

In the vector form

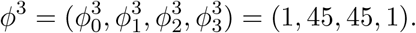

##### Network T4

We consider states *E*_0_ = (0, 0, 0, 0), *E*_1_ = (1, 0, 0, 0), *E*_2_ = (1, 1, 0, 0), *E*_3_ = (1, 1, 1, 0), *E*_4_ = (1, 1, 1, 1). The numbers 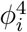 can be computed as above from *u*^4^(*j*), *v*^4^(*j*) and give

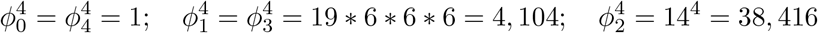

which in the vector form is

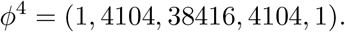

##### Network T5

We consider states *E*_0_, *E*_1_, *E*_2_, *E*_3_, *E*_4_, *E*_5_. The numbers 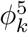 are

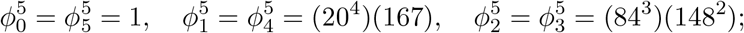

and in the vector form

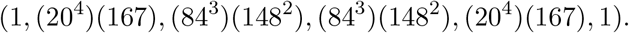

##### Network T6

We consider states *E*_0_, *E*_1_, *E*_2_, *E*_3_, *E*_4_, *E*_5_, *E*_6_. The numbers 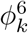 are

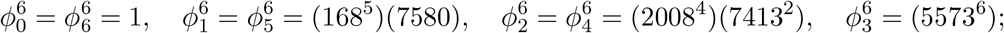

and in the vector form

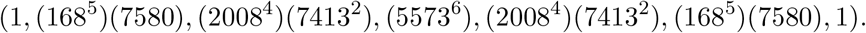

## Data and code availability

The codes used for simulations, scripts for analysis and the simulated data are available at: https://github.com/Harshavardhan-BV/StemCellGRN

## Author contributions

H.B.V., H.S.B., S.A., K.H. performed research. T.G. and M.K.J. conceptualized research. T.G., M.K.J. and H.L. supervised research. All authors contributed to data analysis, writing and editing the manuscript.

## Acknowledgements

H.B.V. is supported by the Prime Minister’s Research Fellowship (PMRF). M.K.J. is supported by Param Hansa Philanthropies. T.G. was partially supported by NSF grant DMS-1951510. K.H. and H.L. were supported by the Center for Theoretical Biological Physics, NSF PHY-2019745.

## Supplementary Material

**Figure S1:**
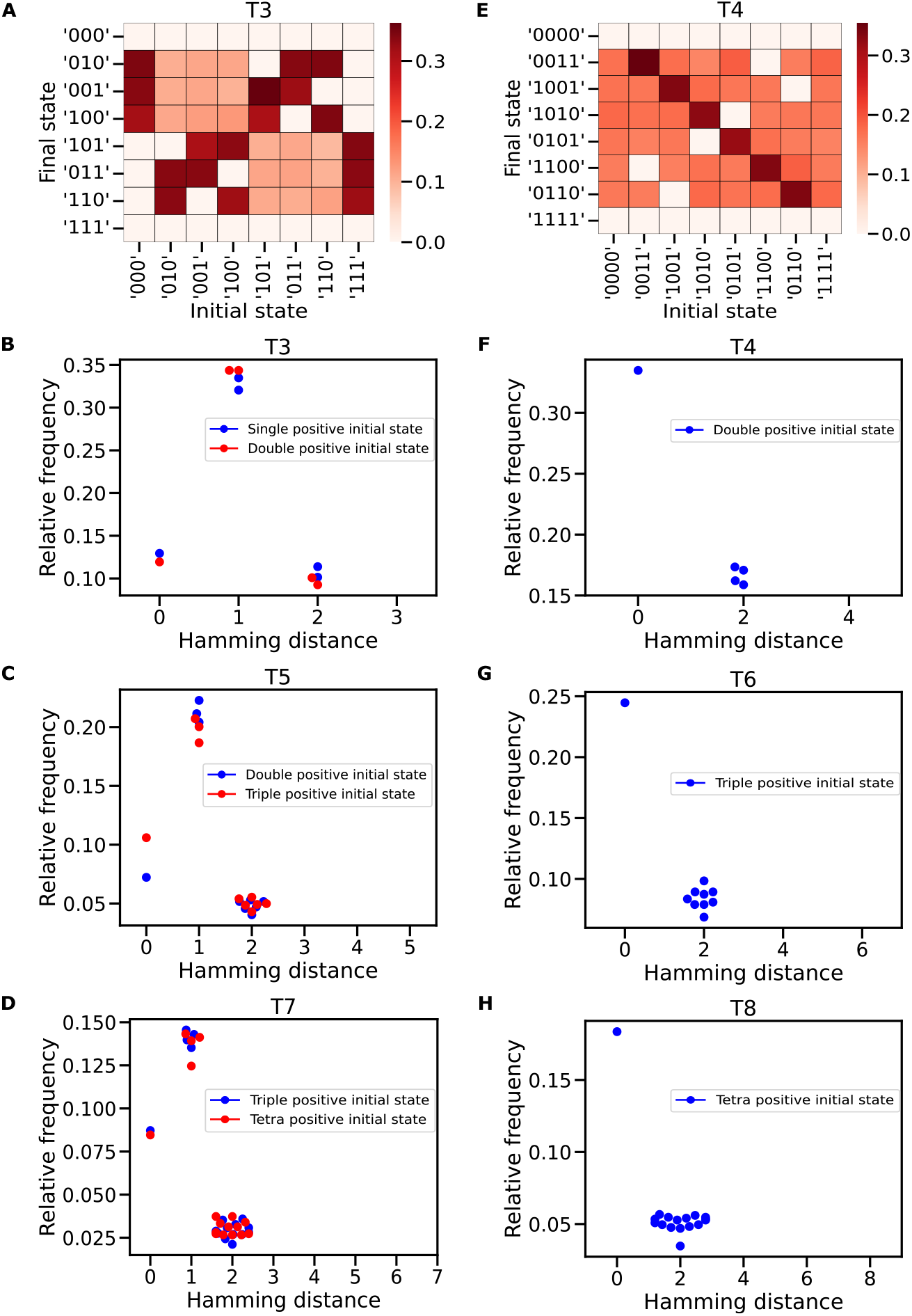
State perturbation dynamics of toggle-n networks: A: Heatmap showing relative frequencies of state transitions on perturbation from a given initial state to a given final state for a representative case of T3. Relative frequencies of state transitions from representative steady states to those steady states at a given hamming distance from the initial state for B: T3, C: T5, and D: T7. E: Heatmap showing relative frequencies of state transitions on perturbation from a given initial state to a given final state for a representative case of T4. Relative frequencies of state transitions from a representative steady state to those steady states at a given hamming distance from the initial state for F: T4, G: T6, and H: T8.

**Figure S2:**
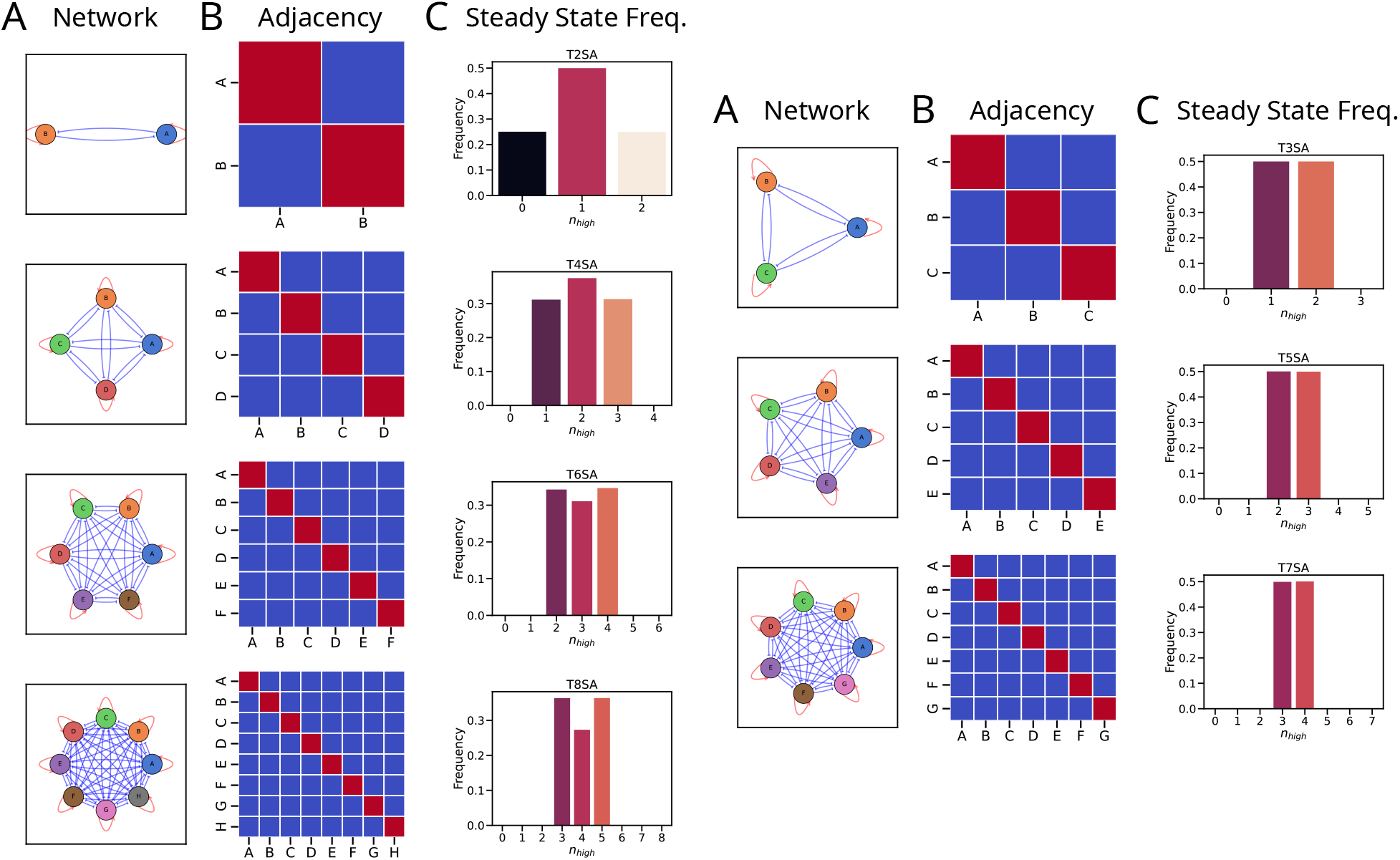
Schematic representation of the toggle-n networks with self-activation and their steady states: A: Graphical network representation illustrating the nodes and edges representing TFs and interactions respectively. B: Adjacency matrix representation with the rows/columns and elements representing the interaction respectively. Red/+1, Blue/-1 and Grey/0 represent activatory, inhibitory and no interactions respectively. C: Steady state frequency distribution of *n*_*high*_ states. Left: Even networks, Right: Odd networks

**Figure S3:**
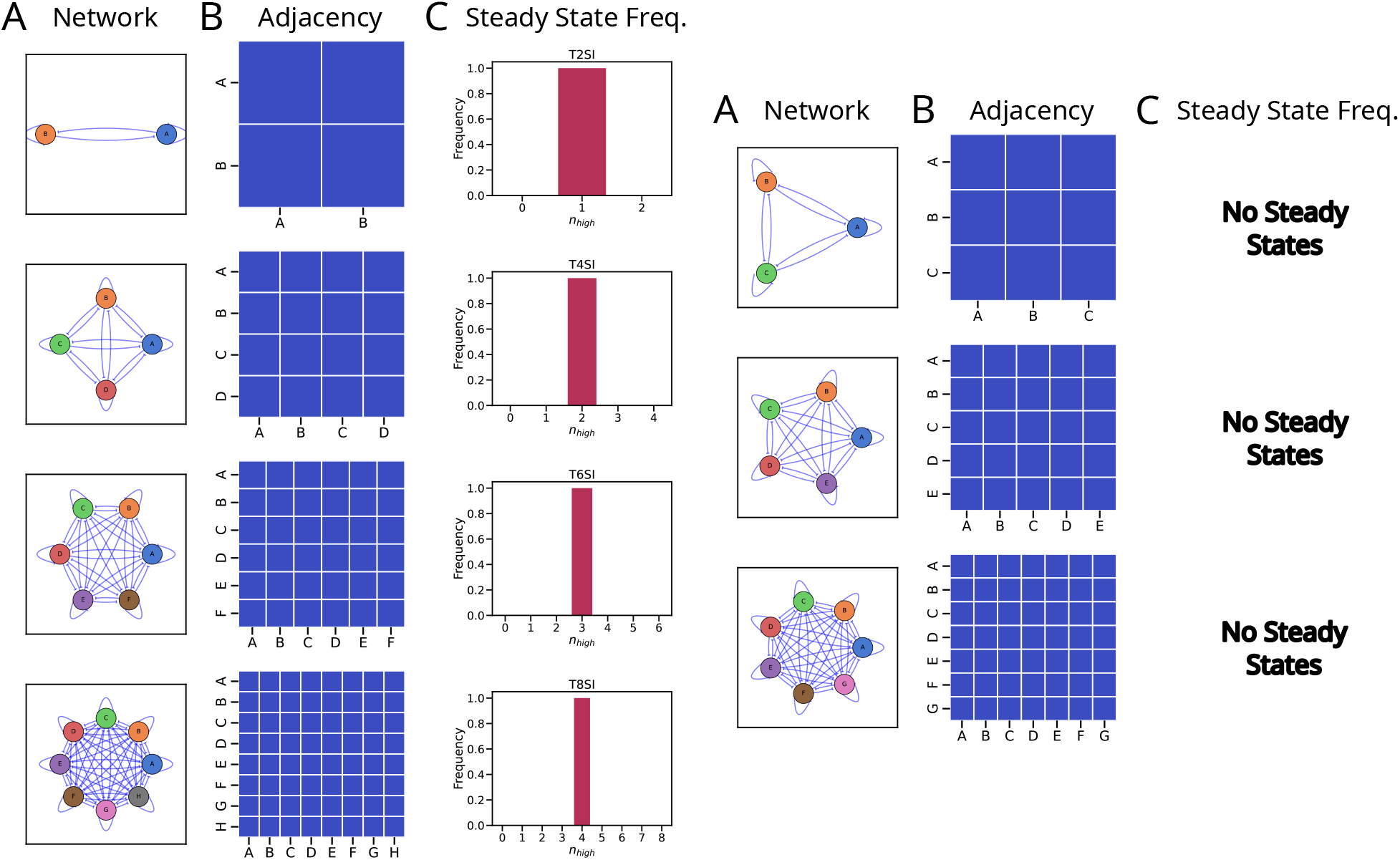
Schematic representation of the toggle-n networks with self-inhibition and their steady states: A: Graphical network representation illustrating the nodes and edges representing TFs and interactions respectively. B: Adjacency matrix representation with the rows/columns and elements representing the interaction respectively. Red/+1, Blue/-1 and Grey/0 represent activatory, inhibitory and no interactions respectively. C: Steady state frequency distribution of *n*_*high*_ states. Left: Even networks, Right: Odd networks

**Figure S4:**
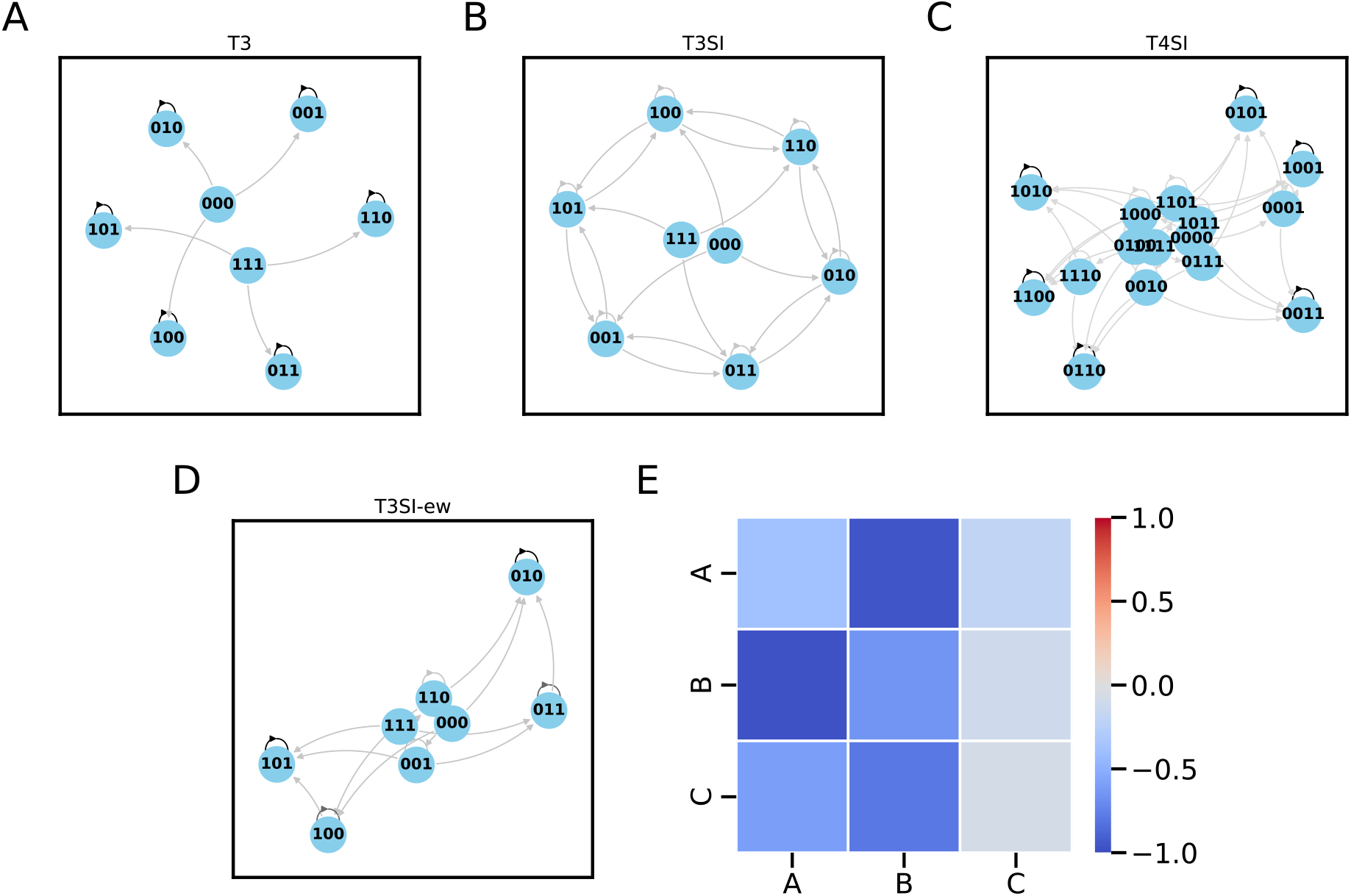
State transition graphs illustrating the stable vs meta-stable states: State transition graphs of A: T3, B: T3 with self-inhibition, C: T4 with self-inhibition, D: T3 with self-inhibition and random edge-weights, E: Shows the corresponding adjacency matrix used for D. The darkness of the edge represents the probability of that transition from the previous state.

**Figure S5:**
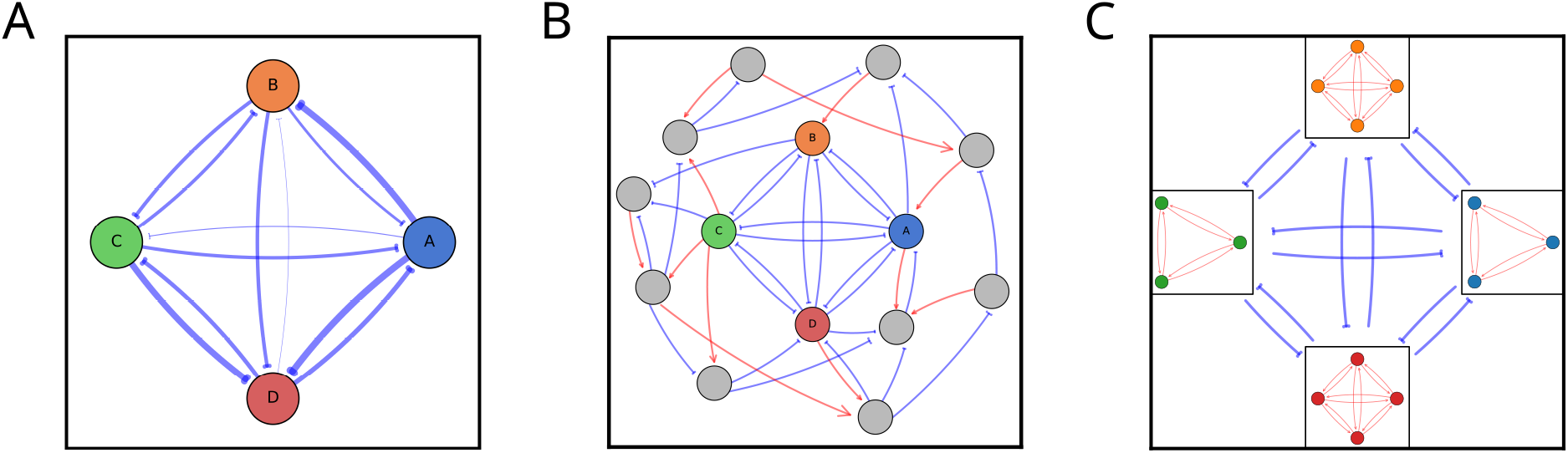
Schematic representations of different perturbations: A: Edge-weight perturbations showing edges with random weights, where edge thickness indicates the strength of regulation. B: Embedding where toggle networks interact with a random network, with grey nodes representing the random network. C: Scaling the toggle-n networks as team-n networks where interactions within teams are activatory, and interactions between teams are inhibitory.

**Figure S6:**
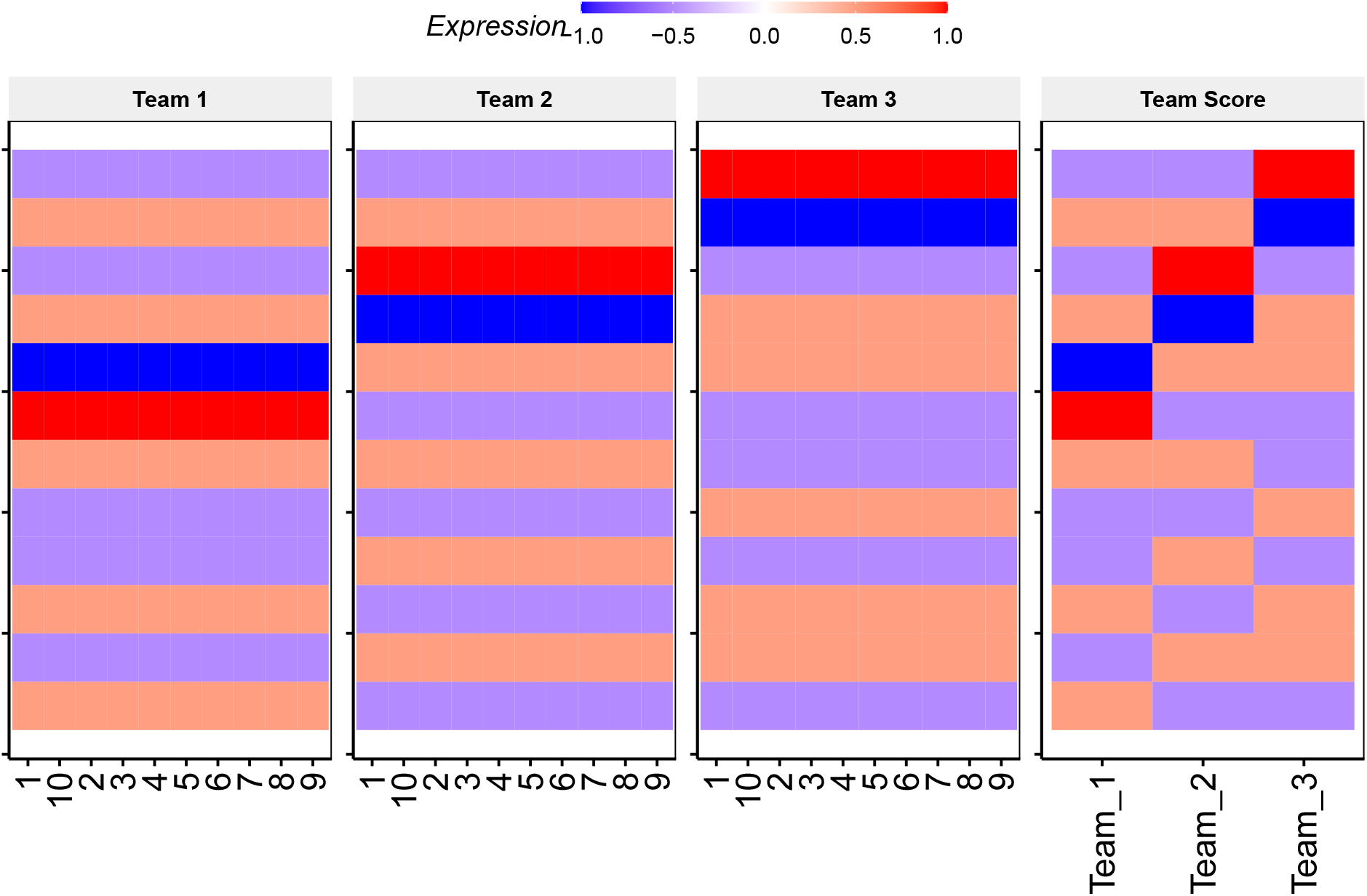
Node expression and team scores for three team network simulated using the four-level model: Expression of nodes grouped by belonging to team 1, team 2, team 3 (left to right) and Team score for each team.

**Figure S7:**
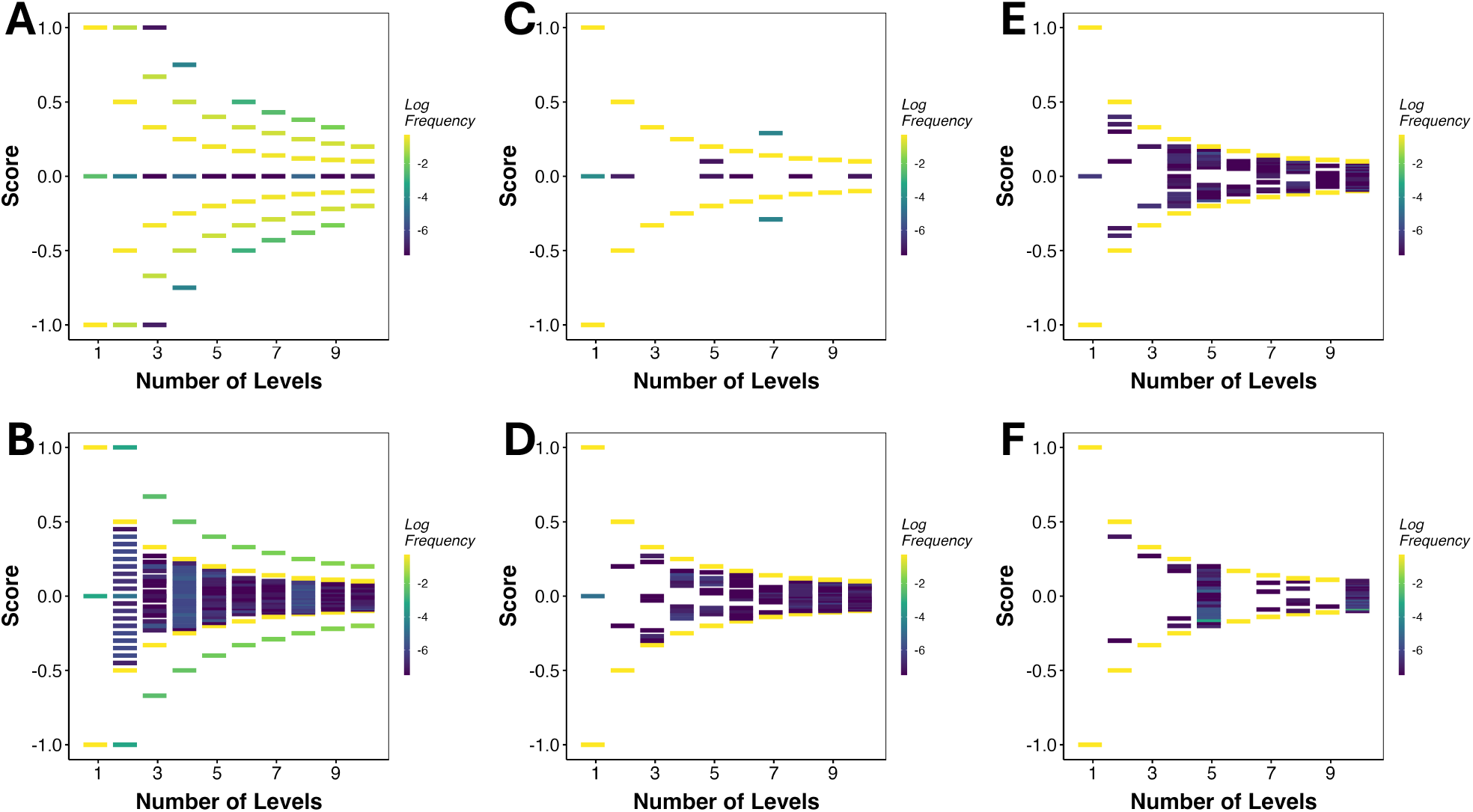
Score of Team 1 across all steady states (y axis) against the number of levels in the multilevel model (x axis): A: three teams, B: four teams, C: five teams, D: six teams, E: eight teams and F: ten team networks.

**Figure S8:**
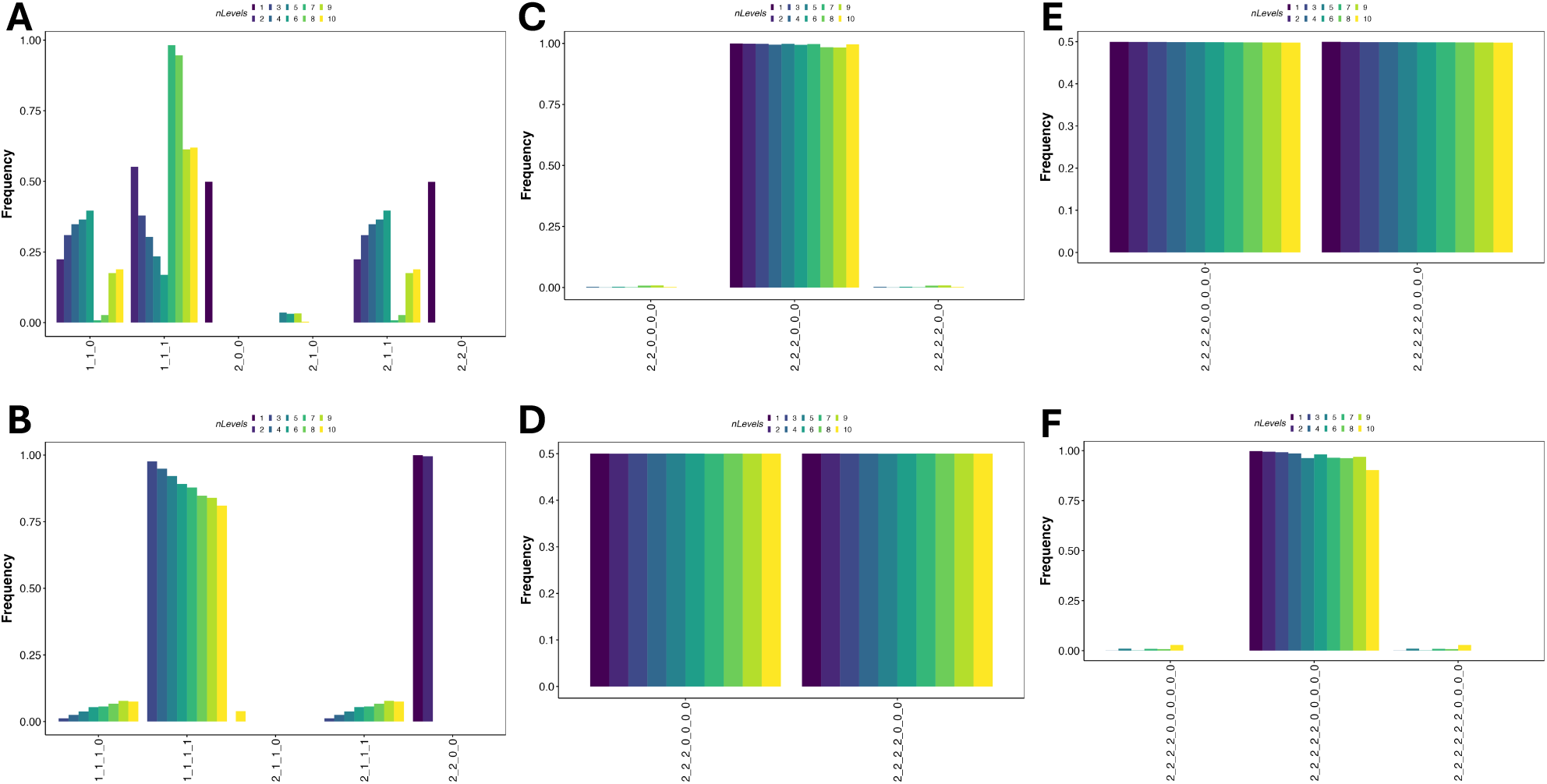
Frequency of normalized and discretized states for networks: A: three teams, B: four teams, C: six teams, D: seven teams, E: nine teams and F: ten teams.

## S.1 Proof for Theorem 1

*Proof*. Let *A* denote the set of nodes with *x*_*i*_ = +1 and *B* denote the set of nodes with *x*_*i*_ = −1. And,

|*A*| = *k* and |*B*| = *s*, with *k* + *s* = *n*

From Font-Clos et al., 2018, the Hamiltonian is defined as:

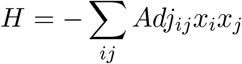

which, for our given system of Tn:

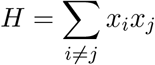

Now the Hamiltonian can be divided into 4 conditions

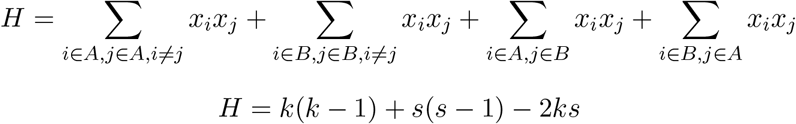

Consider the case when a node flips from *B* → *A*. So, *k*^′^ = *k* + 1 and *s*^′^ = *s* − 1. The Hamiltonian for this case can be written in terms of the original *k* and *s*.

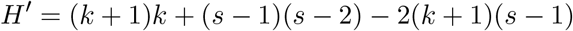

The “energy” change for this new configuration, denoted by Δ*H*^+^:

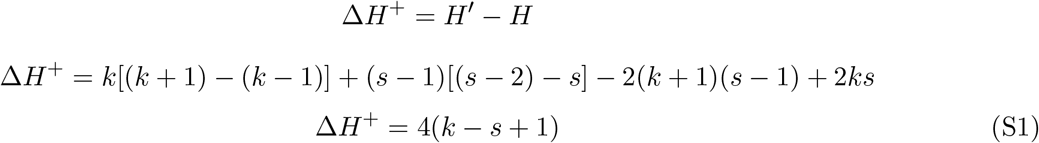

Therefore when *k < s* and we flip vertex from the group of size *s* to a group of size *k* then Δ*H*^+^ *<* 0 and the energy strictly decreases. By the symmetric argument exchanging the identity of groups *A* and *B* it follows that moving an element from larger group to a smaller group decreases the energy (Δ*H*^−^ *<* 0).

We discuss identity of lowest energy states. If the state is in the lowest energy configuration, we must have both Δ*H*^+^ ≥ 0 and Δ*H*^−^ ≥ 0.

Consider the case when *s* = *n* − *k*. Then the condition for Δ*H*^−^ ≥ 0 is 2*n* − *k* + 1 ≥ 0 which gives

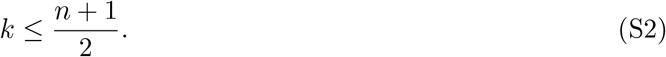

On the other hand the condition for Δ*H*^+^ ≥ 0 reads

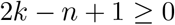

which leads to the condition

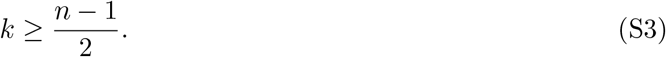

The equations (S2) and (S3) give a constraint for lowest energy states

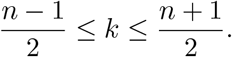

Note that when *n* = 2*k* + 1 is odd, this has solutions *s* = *k* + 1 and *s* = *k*, while when *n* is even the only solution is 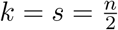. This proves the Theorem.

